# Developing and Characterizing a Murine Model of *In Utero* Transmission of Ebola Virus

**DOI:** 10.64898/2026.08.04.742335

**Authors:** Hanora Van Ert, Corey W. Henderson, Brian J. Smith, Paige T. Donovan, Ethan Kardin, Emma Eubank, Maryam Fakhimi, Matthew Liebermann, Kelly N. Messingham, Mark K. Santillan, Mark L. Schultz, Andrea Marzi, Wendy Maury

## Abstract

Ebola virus (EBOV) disease (EVD) is a hemorrhagic disease caused by EBOV infection. EVD outcomes in pregnant women are similar to non-pregnant women however, EVD is associated with negative fetal outcomes in ∼99% of cases. There is a critical need for a tractable small animal model to study maternal/fetal transmission of EBOV. We utilized interferon α/β receptor knock out mice infected and authentic EBOV or the model virus, recombinant vesicular stomatitis virus encoding EBOV glycoprotein (rVSV/EBOV). Infection with either virus during late pregnancy resulted in placental infection and vertical transmission to the fetus within 2-3 days. Robust levels of maternal and fetal proinflammatory cytokines were evident by day 5 after EBOV infection. Within the placenta, trophoblasts and endothelial cells were viral antigen positive. Elimination of the endosomal receptor NPC1 in junctional zone trophoblasts reduced placental infection and virus transmission to the fetus. These studies establish an infectious model that provides EBOV trafficking and pathogenesis insights during pregnancy.

**Teaser:** This model provides key insights into how viral trafficking and maternal immune responses drive adverse fetal outcomes during gestational Ebola virus infection.

## Introduction

Ebola virus (EBOV) is an enveloped, negative sense RNA virus in the *Filoviridae* family that is associated with significant morbidity and mortality (*1*, *2*). EBOV is a member of the *Orthoebolavirus* genus, along with Sudan virus (SUDV), Bundibugyo virus (BDBV), Tai Forest virus (TFV) and Reston virus (RESV) (*2*). EBOV is a zoonotic virus that once spillover occurs in the human population can also be transmitted person-to-person. Ebola virus disease (EVD) is characterized by systemic viremia, triggering robust inflammatory host cytokine responses, consumptive coagulopathy, hemorrhage, and massive fluid loss leading to multi-organ failure in its victims (*1*, *3*, *4*). Case fatality rates are variable and range between 28-90% depending on the outbreak (*1*, *4*, *5*). In EBOV endemic regions, women who principally serve in care-taking roles are at an increased risk of EBOV exposure compared to men (*6–8*).

EBOV gains access to the host by breaching an epithelial barrier such as the skin or mucosal surface (*2*, *9*, *10*). EBOV enters cells by interactions with a number of different surface attachment factors and ultimately accessing the endocytic pathway. The viral glycoprotein (GP) is proteolytically processed by low pH proteases within the late endosome and the cleaved GP binds to its receptor, the late endosomal/lysosomal host cell protein Niemann-Pick Type C1 (NPC1) (*11*, *12*). Initial viral targets are thought to be mononuclear phagocytes such as macrophages and dendritic cells (DCs) at or near the site of infection (*13–16*). Active EBOV replication in these cells generates infectious virus while inhibiting host innate interferon responses (*17–19*). Virus disseminates to secondary lymphoid organs and as the infection progresses EBOV is found in all major organs in a wide range of cell types including myeloid, stromal, epithelial, and endothelial populations (*16*, *20*). EBOV pathology stems both from direct tissue damage caused by viral infection of tissues as well as coagulation dysfunction and robust cytokine and chemokine responses resulting in a cytokine storm (*1*, *3*).

Data on EVD during pregnancy is limited. However, recent systemic reviews and meta-analyses indicate that case fatality ratios for pregnant women are roughly 68-72% which suggest that pregnant women do not have significantly elevated odds-ratio of succumbing to EVD compared with non-pregnant women (*21*, *22*). In contrast, fetuses and neonates experience devastating outcomes (*21*, *22*), with loss of the fetus occurring in ∼77% of cases and the loss of neonates in ∼99% of births (*22*). The cause of the profound fetal/neonatal loss is not clear but likely includes a combination of direct viral infection of maternal and fetal tissues, coupled with cytokine and physiologic responses of mother and fetus. Further, these data highlight profound loss of pregnancies and neonates that may not be attributable to maternal death. While these data reflect the best information that is available, EVD in pregnancy remains woefully understudied. Barriers to information include the sporadic nature of EVD outbreaks in conjunction with settings that limit the resources needed to adequately document cases. Additionally, variable case report definitions, inconsistent viral testing/isolation protocols, patient reporting, and presentation biases influence data availability and reliability (*21–25*). The combination of these factors results in very small data sets, which make meaningful interpretations difficult and limit knowledge regarding care for pregnant women and their offspring during EBOV outbreaks.

Available filovirus-infected human samples suggest that vertical transmission of EBOV occurs *in utero* across the placenta. Immunohistochemistry (IHC) staining for viral antigen in two placentas from filovirus-infected women demonstrated antigen positivity in villous tissues (*26*). Based on the location and morphology of the cells, the viral antigen-positive cells were likely syncytiotrophoblasts (STBs), cytotrophoblasts (CTBs), and maternal myeloid cells within the intervillous space. In addition, case reports indicate that 23/24 (96%) of tissue samples from infants born to EVD-positive mothers are positive for viral RNA (*22*). Consistent with vertical transmission, a study of experimentally EBOV-infected Angolan free-tailed bats demonstrated *in utero* fetal infection following maternal systemic infection (*27*).

Here, we establish a tractable murine model of EBOV transmission and pathogenesis during pregnancy. Using our model, we define the timing of viral transmission to fetuses during late gestation. Additionally, we identify cells within placental and fetal tissues that support virus infection. Conditional knockout of the endosomal EBOV receptor NPC1, in junctional zone trophoblasts reduced infectious virus in the placentas and transmission of virus to the fetus. EBOV infection results in robust proinflammatory responses by maternal and fetal tissues that likely contribute to fetal demise. Future utilization of this model will provide additional mechanistic insights into *in utero* filovirus transmission, thereby improving obstetric care during EBOV outbreaks.

## Results

### Establishment of a murine model of intraperitoneal (i.p.) rVSV/EBOV infection during late pregnancy

Initial studies to establish our pregnancy model were performed using a BSL2 EBOV infection model based on recombinant vesicular stomatitis virus (rVSV) encoding the EBOV GP gene and GFP reporter in place of the native G gene (rVSV/EBOV). rVSV/EBOV has a similar cell tropism as EBOV, allowing us to define parameters important for EBOV maternal/fetal transmission in BSL2 conditions (*11*, *12*, *28–32*). The use of whole-body homozygous interferon α/β receptor deficient (*Ifnar^-/-^*) mice was required to facilitate systemic infection in this rVSV-based model (*9*, *33*). Dams were infected in late gestation, on embryonic day 14 (E14) to maximize placental maturity and functionality in both nutrient provision and anti-microbial defense (*34*, *35*). These initial studies used intraperitoneal (i.p.) delivery of virus since we sought to model EBOV infection, and this delivery route is the only route that causes systemic infection and pathology following infection of immunocompetent mice with mouse-adapted EBOV (*14*).

Intraperitoneal infection with rVSV/EBOV of pregnant versus non-pregnant females resulted in similar pathogenesis **(Fig. 1A).** To assess if viral replication was elevated in tissues of pregnant dams, titers were performed and we did not observe differences between the infected pregnant and non-pregnant females at 3 days post infection (DPI), a timepoint during rVSV/EBOV infection when high levels of virus can be readily detected in a variety of visceral organs and serum (**Fig. S1A-B**) (*9*, *30*). These data suggest that pregnancy does not pre-dispose to higher infectious titer production, nor outcomes during rVSV/EBOV infection.

**Fig. 1:**
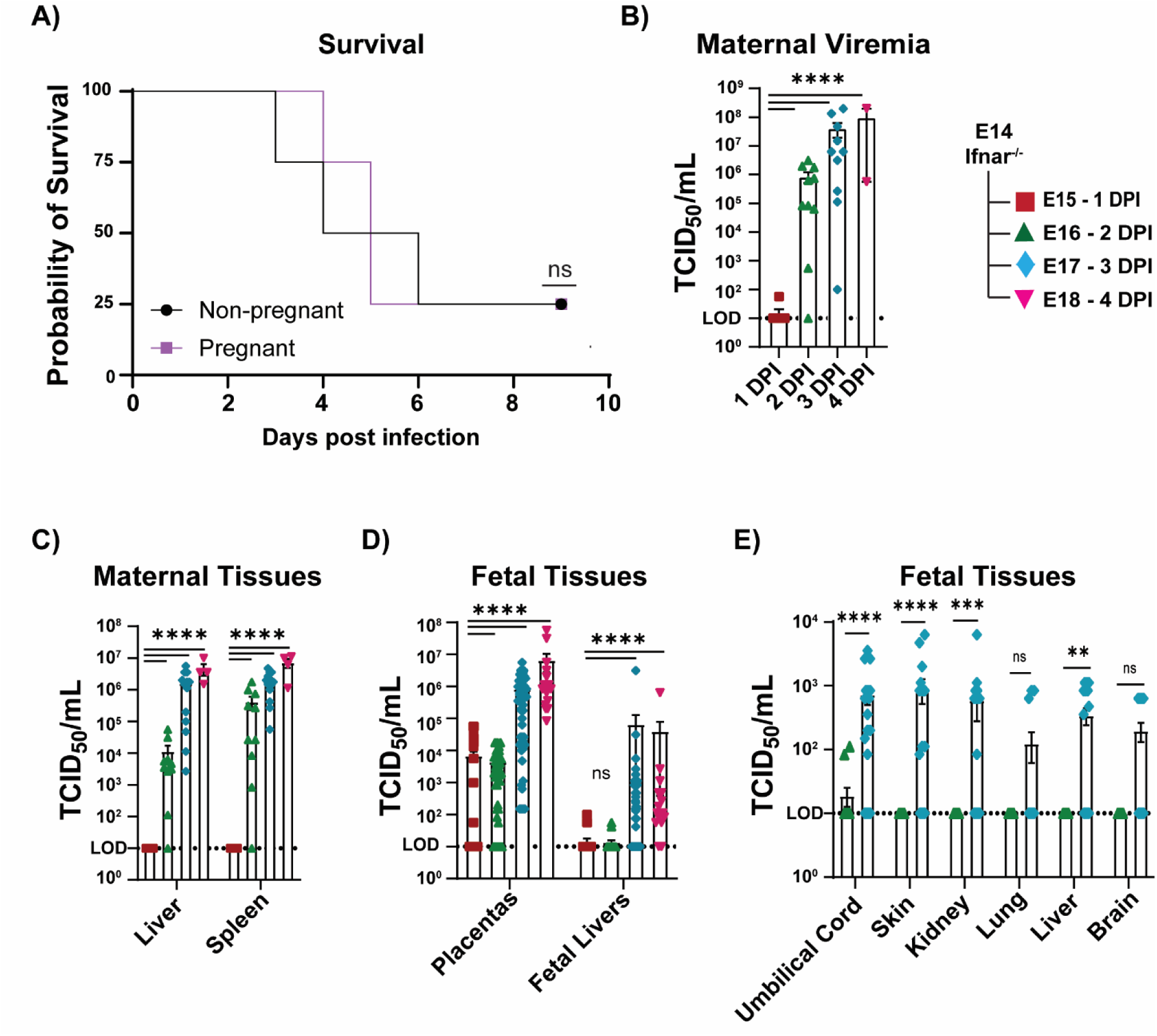
rVSVIEBOV administered i.p. during late gestation mediates robust infection and vertical tranmission. A) Survival is equivalent in E14 *lfnar^-/-^·* dams or age-matched female *lfnar^-/-^* mice infected i.p. with 500 TCID_50_ of rVSV/EBOV. **B-D)** E14 *lfnar^-/-^* dams were infected i.p. with rVSV/EBOV and maternal serum **(B),** maternal visceral tissues **(C)** or fetal tissues **(D)** were harvested at denoted time points and viral titers were quantified (1 DPI n = 9 dams, 2 DPI n = 10 dams, 3 DPI n = 13 dams, and 4 DPI n = 4). Viral titers were determined by TCID_50_ assays on Vero E6 cells. E) Fetal tissues from *lfnar^-/-^·* dams infected i.p. at E14 and harvested at 2 or 3 DPI (n = 5 dams/day). ns = not significant, *p≤0.05, **p≤0.01, ***p≤0.001, ****p≤0.0001 as determined by **(A)** Mantel-Cox test, **(B)** One-way ANOVA w/ Dunnett’s multiple comparisons, **(C-E)** two-way ANOVA w/ Sidak’s multiple comparisons test.

rVSV/EBOV titers in maternal and fetal tissues were evaluated on days 1-4 of an E14 infection of *Ifnar^-/-^* dams (**Fig. 1B-E**). Viremia was detectable in one dam as early as 1 day post infection (DPI) and titers in maternal serum, liver, and spleen increased significantly across time (**Fig. 1B-C**). Most placentas had detectible titers as early as 1 DPI that significantly increased over time, with 100% of placentas containing infectious virus by 3DPI (**Fig. 1D, Fig. S1G**). Titers in fetal liver lagged behind those found in the dams and in placentas with significant titers only evident starting at 3DPI. High viral loads were also detected by RT-qPCR in maternal and placental tissue by 3 DPI (**Fig. S1C**) and positively correlated with infectious titers, especially in placental tissues (**Fig. S1D and E**). In another set of studies, viral titers were assessed in a variety of different fetal tissues, with significant infection observed in most tissues by 3DPI **(Fig. 1E)**. Transmission frequency (percentage of infected tissues relative to the total number of fetuses analyzed) also demonstrated that infectious virus in fetal tissues was delayed compared to maternal and placental tissues (**Fig. S1G**).

As our low containment infection model was composed of the rVSV backbone, we compared rVSV/EBOV and rVSV/G i.p. infections of E14 *Ifnar^-/-^*dams at 3 DPI. While no difference in maternal viremia was evident between dams infected with rVSV/G vs rVSV/EBOV (**Fig. S2A**), significantly lower rVSV/G titers were found in maternal and placental tissues and no rVSV/G was detected in fetal livers on 3DPI (**Fig. S2B-C, Table 1**). These data demonstrate that EBOV GP-driven tropism facilitates placental infection and vertical transmission in pregnant mice.

**Table 1:**
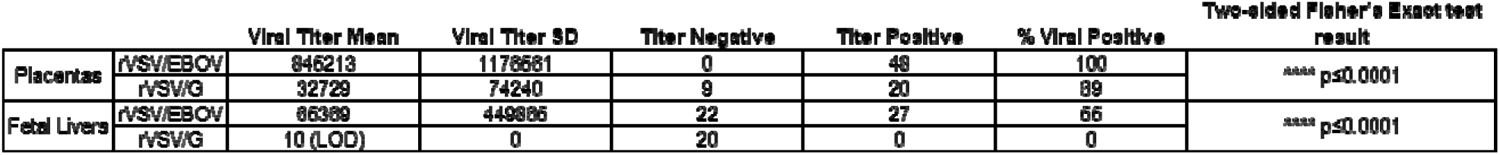
EBOV GP facilitates vertical transmission. Detection of rVSV/EBOV vs rVSV/G in placental and fetal liver tissues at 3 DPI. Dams were infected with 500 TCID_50_ of rVSV/EBOV or 500 TCID_50_ rVSV/G on E14. For each tissue, titer mean and standard deviation were calculated. Additionally, tissues were divided by whether they produced infectious viral titer or not. ****p≤0.0001 two- sided Fisher’s Exact test was calculated on these categorial data. LOD = limit of detection.

Some E14 rVSV/EBOV-infected dams survived infection to parturition allowing us to evaluate infection on parturition timelines with decreasing doses of viral inoculum (**Fig. S3A-B**). Infected dams experienced significantly earlier parturition than PBS injected dams, with 75% of dams given a LD_75_ dose delivering by E17 or E18 (**Fig. S3B**). No difference in parturition was seen in dams given a LD_25_ dose versus PBS. rVSV/EBOV infection on E14 did not alter E17 placental weights (**Fig. S3C**). However, there was a slight reduction (∼8%) in fetal weights and placental efficiency (ratio of fetal and placental weight) (**Fig. S3D-E**) (*36*). As some fetuses from rVSV/EBOV-infected dams did not have detectable infectious virus within their livers at 3 DPI, (**Fig. 1D**) we compared weights of fetuses whose livers did or did not contain infectious virus and these weights did not differ significantly (**Fig. S3F**). Gross morphological abnormalities were also not observed in pups harvested from rVSV/EBOV infected dams (**Fig. S3G**) and the numbers of total concepti per dam (inclusive of resorptions and live pups) were not statistically different (**Fig. S3H**). *In utero* demise of infected fetuses was also not observed (**Fig. S3I**).

### rVSV/EBOV infection of E5 *Ifnar^-/-^* dams resulted in infected concepti tissues with reduced concepti weights

To examine the impact of rVSV/EBOV i.p. infection of *Ifnar^-/-^* dams at earlier stages of pregnancy, studies were also conducted at E5, with maternal and embryonic tissues harvested at E8. Separation of the embryo from developing placental structures was not possible with these early concepti, so whole conceptus tissue was assessed for viral titers. Similar to late gestation infections, high viral titers (10^3^ to >10^8^) in maternal samples and concepti were observed (**Fig. S4A**). A modest reduction of infected concepti weights was observed, suggesting that infection may disrupt placental and/or fetal development during this crucial developmental window (**Fig. S4B**).

### Intravenous (i.v.) and intramuscular (i.m.) rVSV/EBOV infection model hematogenous vertical transmission during late gestation

Our i.p. studies demonstrated robust infection in placentas at 1 DPI (**Fig. 1D**). At this time, minimal to no viral replication was evident in maternal tissues, suggesting direct infection of uterine tissue from the peritoneum occurred (**Fig. 1C-D**). Since direct viral transmission from the peritoneum to fetal tissues does not represent a hematogenous/systemic route of infection from the dam to the fetus, we assessed i.v. and i.m. routes of delivery. E14 *Ifnar^-/-^*dams were infected and maternal, placental, and fetal tissues were harvested at 1-4 DPI (E15-18 of gestation). We administered a viral dose of 5×10^6^ TCID_50_ which is reflective of viremic titers seen in our i.p.-infected dams on 2-3 DPI (**Fig. 1B**). With i.v. delivery, maternal viremia was evident at 1 DPI (**Fig. 2A**), accompanied by appreciable liver and splenic infection (**Fig. 2B**). Maternal serum and liver titers significantly increased across the timespan of infection, while maternal spleen titers were high at 1 DPI and did not increase over time (**Fig. 2B**), suggesting a homing of infectious virus to this organ during this route of infection. The majority of placentas contained infectious virus at 1 DPI; however, titers were 1-4 logs lower than those of maternal titers at this time, indicating maternal systemic infection preceded placental infection and hematogenous spread of the virus to the placenta (**Fig. 2C**). Infection of the fetuses lagged behind maternal and placental tissues, showing minimal viral replication until 3 DPI (**Fig. 2C**). In a second cohort of E14 infected dams, virus infection of E17 fetal tissues was evaluated (**Fig. 2D**). While infectious virus was present in greater than 80% of umbilical cord and skin tissue, 55-70% of visceral organs and brain had detectable virus, suggesting that virus reaches those fetal organs more slowly. To independently verify that cells within fetal tissues were infected and that titers were not due to maternal blood contamination, fetal liver was immunostained for viral antigen (GFP) as well as the macrophage marker, F4/80. F4/80 positive cells within fetal livers from i.v. inoculated dams co-stained for viral antigen, highlighting fetal liver macrophages as a potentially relevant cell type supporting viral infection (**Fig. 2E**). Fetal growth parameters were also assessed at 3 DPI. We observed no significant difference between infected and uninfected samples in the placental or fetal weights, placental efficiency, number of total concepti per pregnancy, or the morphology of fetuses (**Fig. S5**).

**Fig. 2:**
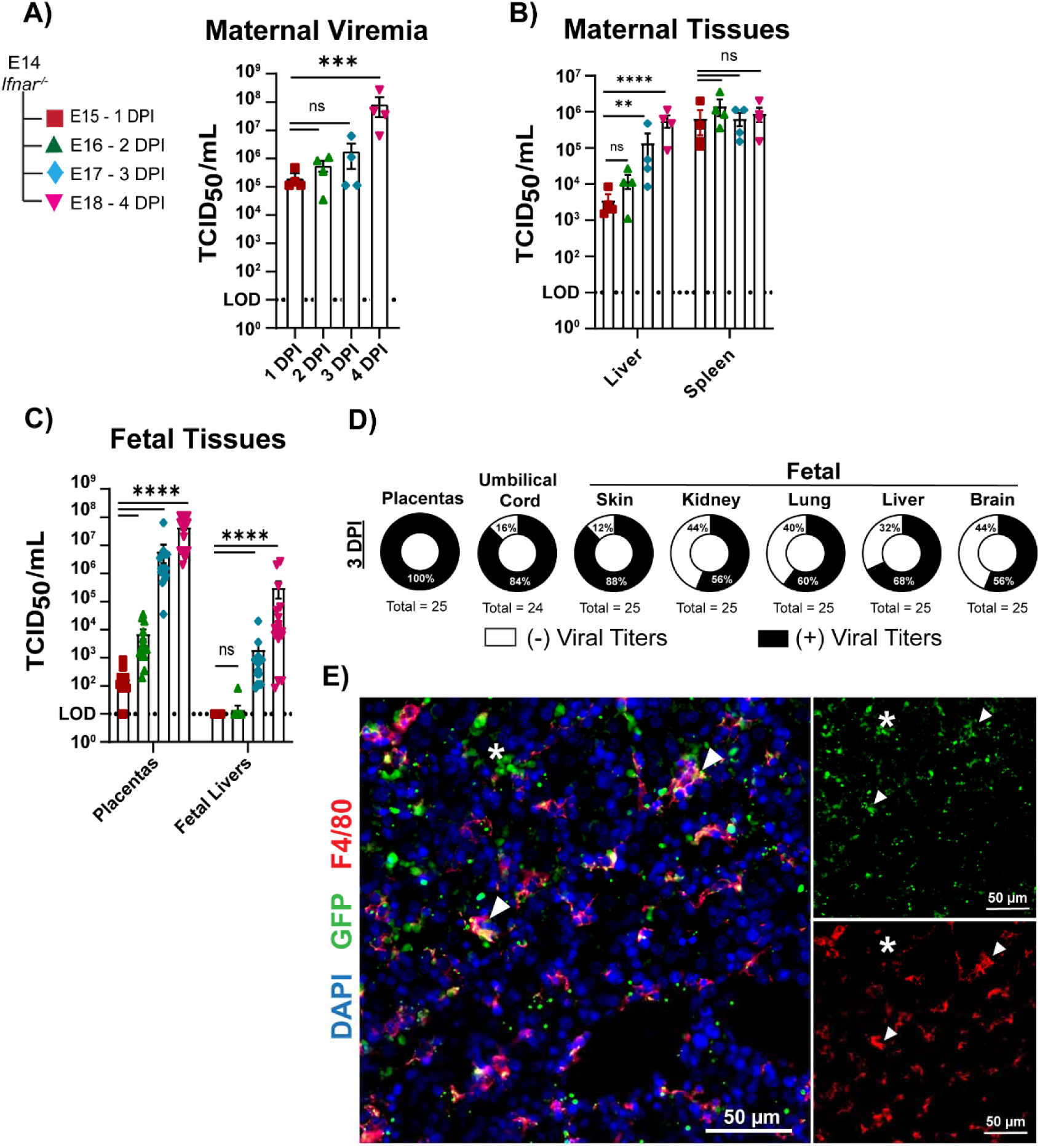
Intravenous rVSV/EBOV infection results in systemic viral dissemination in maternal, placental, and fetal tissues. **A-C)** E14 */fnar^-/-^* dams were infected with 5×10^6^TCID_50_ of rVSV/EBOV i.v. and **(A)** maternal serum, **(B)** maternal visceral tissues or **(C)** fetal tissues were harvested at denoted timepoints post infection. Titers were quantified via TCID_50_ assays on Vero E6 cells. n **4** dams per day. **(D)** Percent of placental and fetal samples with detectable viral titers in an independent cohort of *lfnar^-/-^* dams. n = 3 dams. **(E)** 3 DPI fetal liver immunostained for GFP and F4/80 antigen. Co-staining marked with arrowheads. Cluster of cells solely staining for viral antigen staining is marked with asterisk. Representative image of n 4 fetuses harvested from n 2 i.v challenged dams. 20x image. Brightness and contrast were adjusted uniformly across the image. ns = not significant. *p≤0.05, **p≤0.01. ***p≤0.001, ****p≤0.0001 as determined by **(A)** one-way ANOVA w/ Dunnett’s multiple comparisons, or **(B-C)** two-way ANOVA w/ Sidak’s multiple comparisons test.

We also assessed i.m., subcutaneous (s.q.) and intravaginal (i.vag.) administration of rVSV/EBOV. While placental and fetal infection via s.q. or i.vag. was not detected (**Fig. S6**), i.m. delivery resulted in similar levels of infection as the i.v. route, with 100% placentas and 60% of fetal livers infected on 3DPI (**Fig. S7A and B**). A prior study demonstrated that an i.m. route of EBOV delivery results in effective systemic infection of non-pregnant females (*37*). With i.m. delivery, we found that placental weights were modestly reduced; however, this difference in placental weights was not observed with fetal weights, resulting in modestly enhanced placental efficiency (**Fig. S7C-E**). There was also no difference in the number of concepti per pregnancy, or fetal morphology between the two treatment groups. (**Fig. S7F-G**).

### Intramuscular EBOV-Mayinga infection of E13 *Ifnar^-/-^* dams resulted in systemic infection and vertical transmission

To determine if authentic EBOV crosses the placental barrier in our mouse model, we infected E13 *Ifnar^-/-^* dams and non-pregnant females with 10^4^ PFU of EBOV (Mayinga) i.m. and tissues were harvested at 3, 4, and 5 DPI (E16-18). We chose to harvest at later days post-infection with EBOV to account for the longer replication period required by authentic EBOV compared to rVSV/EBOV. Initial investigations compared viral loads and infectious titers in pregnant and non-pregnant females at 5 DPI (**Fig. S8**). Viral loads trended higher in tissues of pregnant dams. However, these trends were not statistically different. In the pregnant dams, high levels of EBOV RNA and infectious virus were found in maternal blood and spleen as early as 3 DPI, and levels did not change at 4 or 5 DPI (**Fig. 3A, B, D and E**). Within the liver, virus load and titers trended upward, but were not statistically significance due to wide variation in values (**Fig. 3B, E**). Similar to maternal tissues, viral RNA and infectious virus were detected within ∼85-95% placentas, with infectious titers increasing across time (**Fig. 3C, F, G**). Viral loads and titers were lower in fetal liver compared to maternal and placental tissues, but trended upward in number of positive samples, and viral load across infection. (**Fig. 3C, F, H**). These data provide experimental evidence of EBOV transmission in utero.

**Fig. 3:**
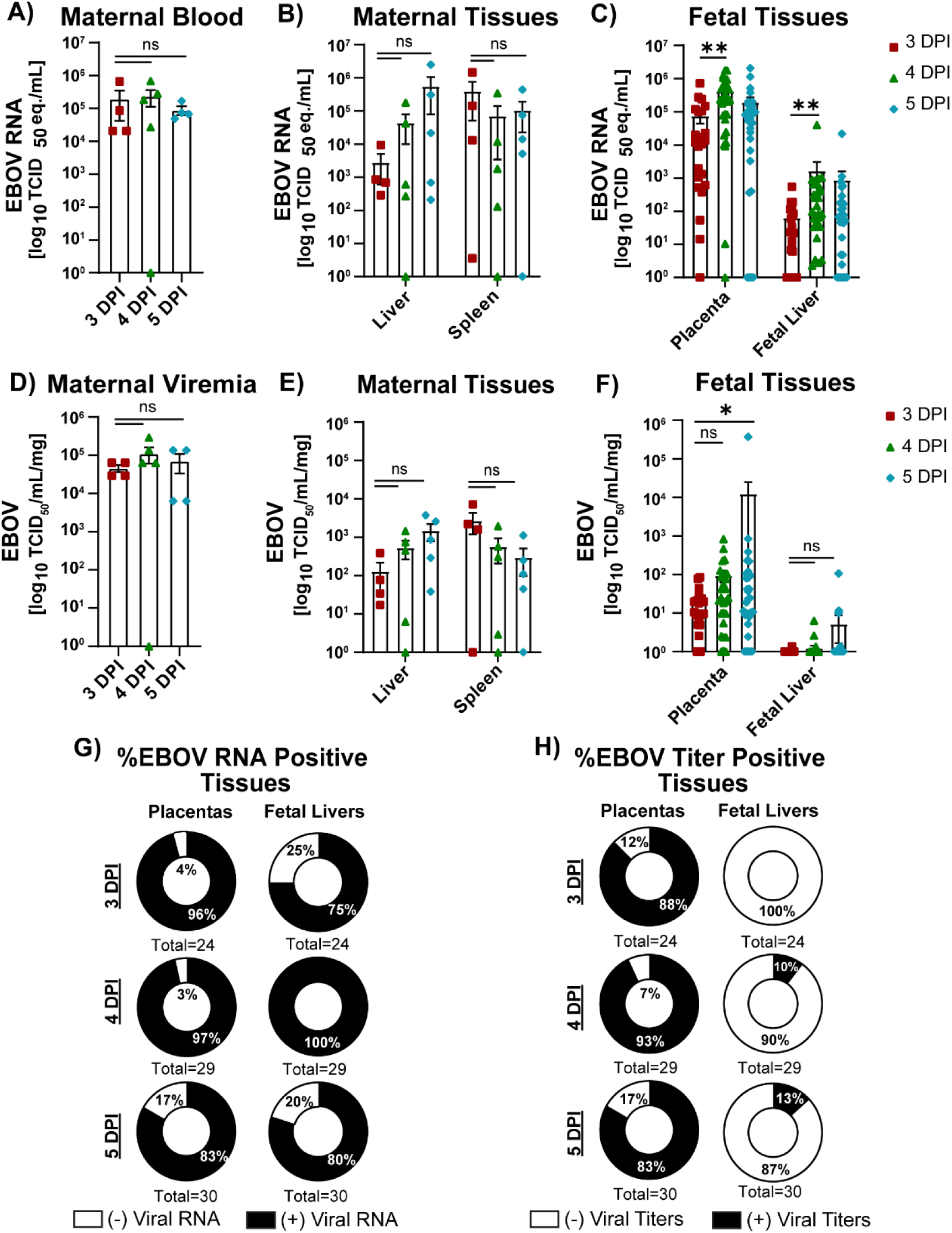
Intramuscular inoculation of EBOV results in vertical transmission in *lfnar^-/-^* mice. E13 *lfnar·* dams were infected with 1o^4^PFU of EBOV-Mayinga i.m. and tissues were harvested at 3 DPI (E16), 4 DPI (E17), and 5 DPI (E18). **A-C)** Viral RNA quantified by RT-qPCR and normalized for tissue weight where relevant from **(A)** maternal blood, **(B)** maternal visceral tissues, and **(C)** placental and fetal livers. **D-F)** Infectious viral titers normalized to harvested tissue weights from **(D)** maternal blood, (E) maternal visceral tissues, and **(F)** placentas and fetal livers. **G-H)** Percent of fetal tissues containing **(G)** viral RNA, or **(H)** viral titers. **N** = 4-5 dams per day, n = 24-30 placentas/fetal livers per day. ns = not significant, *p:≤0.05, **p :≤0.01, ***p:≤0.001, ****p:≤0.0001 as determined by (A, D) one-way ANOVA w/ Dunnett’s multiple comparisons, or (B-C, E-F) two-way ANOVA w/ Sidak’s multiple comparisons test.

### EBOV antigen localized predominantly to trophoblasts

Murine placentas contain both maternal- and fetal-derived components (*34*). Maternal tissue comprises the minority of the structure and is termed the decidua (**Fig. 4A**). Much of the murine fetal-derived placental is comprised of a variety of different trophoblasts that are abundant within the junctional zone (JZ) and the labyrinth zone (LZ) (*34*, *38*, *39*). These include the spongiotrophoblasts, glycogen cells and parietal giant trophoblasts in the JZ and syncytiotrophoblasts and sinusoidal giant trophoblasts in the LZ (*39–41*). Endothelial cells and macrophages also serve as crucial placental cell types and are known to be targets of EBOV in other tissues (*42–44*). To evaluate which cell types within the placenta support virus infection, placentas from EBOV-challenged dams were immunostained for EBOV glycoprotein (GP) and we found all regions of the placenta contained detectable viral antigen. However, cells within the decidua/junctional zone regions had the most robust EBOV GP staining (**Fig. 4B, arrowhead**), with viral antigen also localized focally within the labyrinth (**Fig. 4B; inset 1**) and deeper in the placenta along the chorionic plate (**Fig. 4B; inset 2**).

**Fig. 4:**
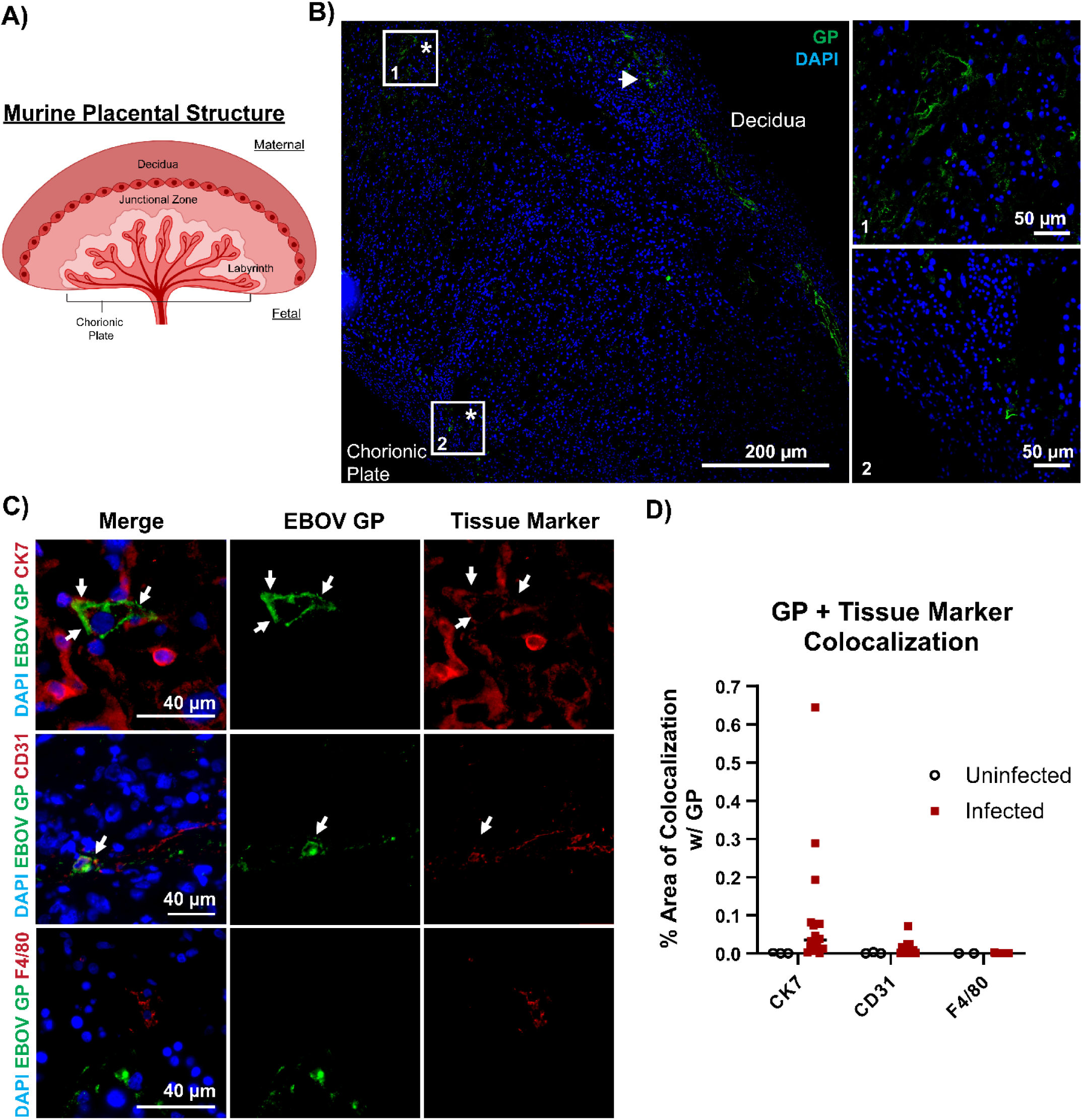
Placental trophoblasts and endothelial cells are infected by EBOV. **A)** Cartoon depiction of murine placental structures. **B-E)** lmmunostaining of 5 DPI placental sections. E13 *lfnar*^-/-^*•* dams were infected i.m. with 1O^4^PFU of EBOV. **(B)** Placental tissues immunostained for EBOV GP and DAPI (20x tilescan). White arrowhead identify EBOV GP antigen-positive cells within the decidual and junctional zone regions. Insets demonstrate EBOV GP focal positivity in 1. the labyrinth, and 2. the chorionic plate of the placenta. **(C)** Placental sections stained for EBOV GP and CK7 (trophoblasts), CD31 (endothelial cells) and F4/80 (macrophages). Arrows identify co-localization of markers. **(D)** Percent area of each image with colocalized pixels of EBOV GP and individual tissue markers. N = 2 placentas/group with 6-10 images from each placenta. 63x images. **(A)** Cartoon created from www.biorender.com.

To identify cell types sustaining EBOV infection, we co-stained 5 DPI (E18) placentas for EBOV-GP and cell-type specific markers (CK7, trophoblasts; CD31, endothelium; F4/80, macrophages). For each co-stained set of antigens, we calculated the area within each image that demonstrated dual positivity of EBOV GP and the cell marker in placentas harvested from uninfected and EBOV infected dams. In co-stained samples, viral antigen positive cells most robustly co-stained with CK7 (**Fig. 4C, D**), implicating trophoblasts as important cell types in vertical transmission. Instances of viral antigen co-staining with CD31^+^ endothelial cells were also observed. Unexpectedly, detectible co-staining of EBOV GP with F4/80 was not observed, suggesting tissue macrophages are not a predominant cell type supporting placental infection.

To extend these cellular tropism studies, 3 DPI (E17) placentas harvested from *Ifnar^-/-^* dams that were challenged with 5×10^6^ TCID_50_ rVSV/EBOV i.v. were immunostained for GFP expressed from the viral genome. Viral antigen was abundant in the decidua and junctional zones with foci of infection found in other locations throughout placental tissue similar to the EBOV challenged mice (**Fig. S9A, asterisks and inset**). Co-staining with the trophoblast marker CK7 in the rVSV/EBOV infected placentas was evident (**Fig. S9A, B white arrows**). Additionally, occasional co-staining of CD31 and GFP could be seen, indicating infection of endothelial cells in placental tissue (**Fig. S9B**). Sporadic incidences of co-staining of macrophage marker Iba1 and GFP were found in contrast to placentas from EBOV-infected dams (**Fig. S9B**). To validate that the GFP staining pattern recapitulated that of EBOV GP, we also immunostained the same placentas from rVSV/EBOV challenged dams with EBOV GP and CK7. We found that EBOV GP is expressed robustly within these tissues and colocalizes with CK7 (**Fig. S9C**). These data indicate that EBOV and the low containment model virus rVSV/EBOV demonstrate tropism for multiple placental cell types, with trophoblasts being the most predominant. Additionally, cells throughout the placenta were infected, demonstrating broad dissemination of the virus.

### EBOV infection resulted in pro-inflammatory cytokine expression in maternal, placental, and fetal tissues

EBOV infection was evident in placental and fetal tissues and thus may result in viral-mediated pathogenesis in these tissues. As EVD is characterized by profound cytokine disturbances, these may in turn be altered by pregnancy or may have deleterious effects on pregnancy outcomes (*1–4*). To date, no assessment of maternal and fetal cytokine profiles in response to EBOV infection has been performed. Sera from naïve and i.m. EBOV infected dams at 3-5 DPI (E16-18), as well as non-pregnant, EBOV-infected females at the same times post infection, were quantified for 13 cytokines and chemokines via bead-based flow cytometric assay. By 5DPI, a broad range of proinflammatory cytokines/chemokines were elevated by EBOV infection, including IFN-γ, TNF-α and IL-6, which have important roles in negative pregnancy outcomes and may be responsible for placental and fetal tissue damage during EBOV infection (*45*, *46*) (**Fig. 5, S10**). Some of these proteins were also elevated as early as 3DPI during infection with IFN-γ, CXCL10, and CCL2 elevated in comparison to naïve pregnant sera. CXCL10 and CCL2 are responsible for recruiting immune cells to sites of inflammation, thus the significant difference in expression early in infection between non-pregnant and pregnant dams may alter the course of immune reaction and pathophysiology (**Fig. S10D-E**) (*47*, *48*). CXCL1 spiked in the infected groups early during infection but returned to naïve levels at 5 DPI. As this chemokine stimulates angiogenesis, future studies assessing the vascularity of EBOV-infected versus uninfected placental tissue is needed (*49*). Although elevated serum levels of IFN-α and IFN-β typically influence pregnancy outcomes, the inability of *Ifnar^-/-^* mice to respond to type I interferon signaling may confound our observations by obscuring viral-mediated pathology (*50–52*). Future studies are needed to examine the effect of these proinflammatory proteins on pregnancy outcomes during EBOV infection.

**Fig. 5:**
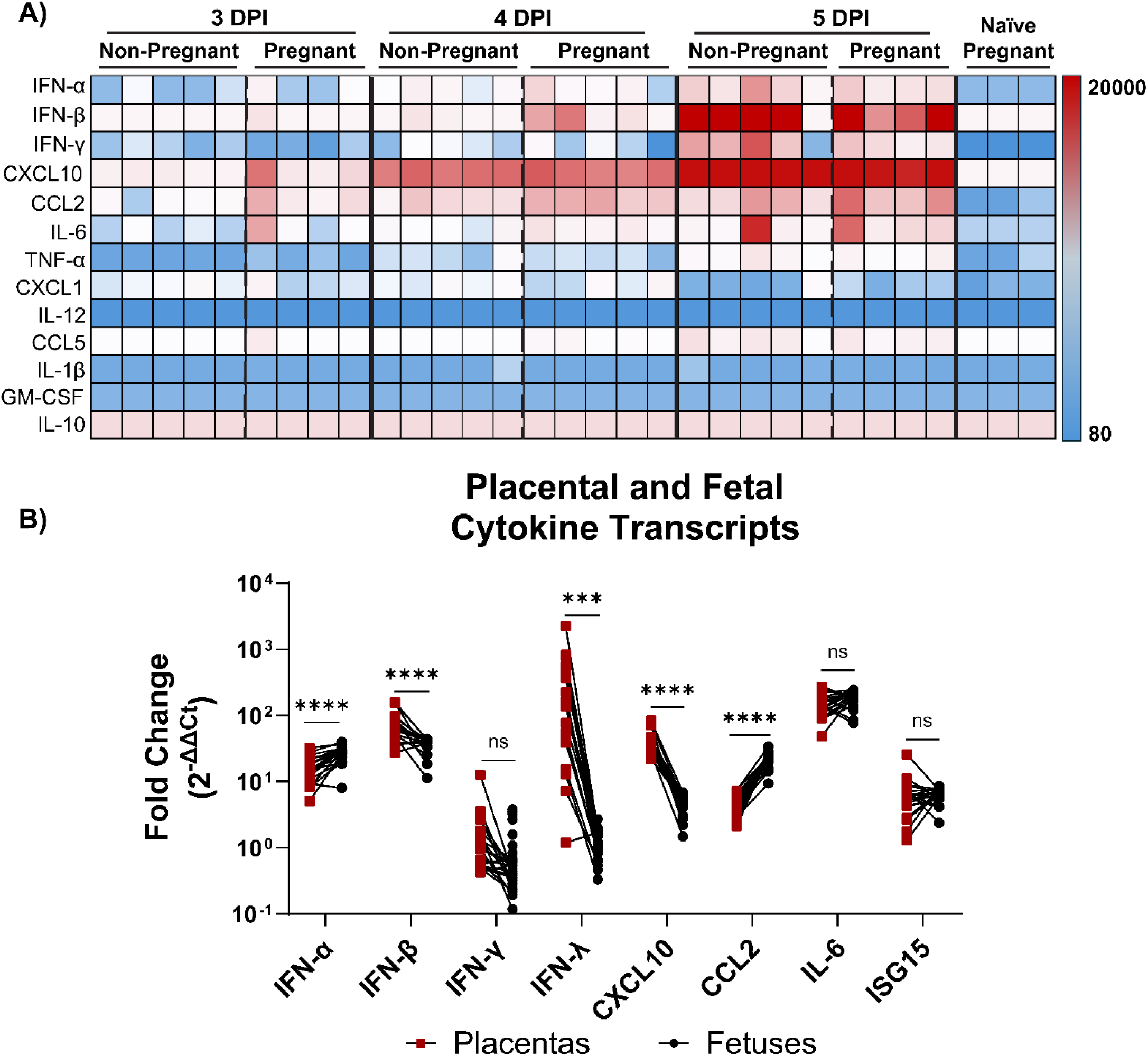
Infection with EBOV results in high expression of inflammatory cytokines and chemokines in maternal sera, placenta and fetal liver. **A)** Cytokine/chemokine profile in *lfnar*^-/-^ dam sera on day 3(E16), day 4 (E17) and day 5 (E18) of infection. E13 dams were infected with 10^4^ PFU of EBOV i.m. Non-pregnant females were infected and harvested in parallel. Serum from uninfected (Naive) pregnant E15-E17 *lfnart*^-/-^ dams were included as negative controls. Scale bar indicates cytokine concentration in pg/ml. Each column represents serum from a single mouse. **B)** Fold increase in RNA expression from placentas and fetal livers at 5 DPI via qPCR. RNA values were normalized to na·1ve-pregnant RNA averages and the fold change was plotted for each placenta-fetus pair. n = 20-30 placenta and fetal liver pairs. ns = not significant, *p≤0.05, **p≤0.01, ***p≤0.001, ****p≤0.0001 as determined (B) paired I-test.

Cytokines and chemokines were measured in serum from 3 DPI of rVSV/EBOV infected dams and uninfected dams (**Fig. S11**). Consistent with the elevated expression of these proteins in EBOV-infected maternal sera, rVSV/EBOV infection stimulated production of proinflammatory proteins, including IFN-γ, IFN-β, CXCL10, CCL5 and IL-6.

Transcript levels of pro-inflammatory genes were also assessed in 5 DPI EBOV-infected placental and fetal tissues and are shown as log_2_-fold expression changes compared to tissues harvested from naïve dams (**Fig. S12**). Except for IFN-γ, all evaluated placental transcripts were elevated in the EBOV-infected tissues, demonstrating a highly proinflammatory placental environment at 5 DPI (**Fig. S12A**). The most notably enhanced transcript in infected placentas was IFN-λ, produced constitutively by placental trophoblasts and enhanced by viral infections (*53*). In EBOV infected placenta, IFN-λ was elevated 500-fold above placentas from uninfected pregnant dams at this late timepoint in pregnancy (**Fig. S12A**). In examination of fetal liver cytokines and chemokines, all transcripts with the exceptions of IFN-γ and IFN-λ were significantly elevated in EBOV-challenged tissues, indicating elevated levels of proinflammatory responses occurring in the tissue (**Fig. S12B**). This upregulation of innate immunity was profound despite relatively modest levels of infectious EBOV evident in fetal tissues (**Fig. 3D, F**). It is possible that this robust innate immune response may facilitate control of viral replication in fetal tissue and needs to be examined further.

A comparison of cytokine/chemokine expressions in infected placenta/fetal pairs was also performed to visualize the similarities and differences in expression between these tissues. We plotted the transcript fold change above naïve tissue for each feto-placental unit (**Fig. 5B**). Trends in innate immune responses were similar across all the pairs analyzed. Paired t-tests highlighted that IFN-γ, ISG15 and IL-6 had similar elevated levels of expression in placentas and fetuses, whereas other cytokines had greater expression in one of the tissues.

### Junctional zone trophoblast-specific NPC1 depletion reduced placental viral titers and fetal transmission

In EBOV and rVSV/EBOV infected placentas, viral antigen co-stained with CK7+ trophoblasts in both the JZ and LZ of the decidua, with the most extensive staining observed within the JZ (**Fig. 4A, Fig. S9**). To examine the importance of trophoblasts in mediating virus placental infection and transmission to fetuses, a Tpbpa/Ada-Cre-driven conditional knockout mouse line was generated that eliminated expression of the EBOV receptor, NPC1, in trophoblastic giant cells and glycogen trophoblasts which are located primarily in the placental JZ (*54*, *55*). On an *Ifnar^-/-^* background, mice expressing the trophoblast-specific enhancer/promoter Tpbpa/Ada-Cre were crossed with NPC1^fl/fl^ mice (*56*, *57*). These mice were found to be healthy and fertile (*58*). The resulting conditional NPC1 knock out mice (Tpbpa/Ada^cre^/Npc1^fl/fl^/*Ifnar^-/-^*(called Ada^cre^/Npc1^fl/fl^)) allowed us to assess the impact of virus infection of JZ trophoblast on placental and fetal viral infection.

To confirm the phenotype of our new mouse line, lysates from *Ifnar^-/-^* and Ada^cre^/Npc1^fl/fl^ E17 placentas were evaluated for NPC1 expression via immunoblots, demonstrating a significant reduction in NPC1 present in the Ada^cre^/Npc1^fl/fl^ placentas (**Fig. S13A**). To verify that Cre was specifically expressed in the trophoblasts within the placenta, trophoblasts were isolated and purified from enzymatically digested placentas followed by density gradient centrifugation (*59*). Cre transcripts were detected by RT-qPCR in the Ada^cre^/Npc1^fl/fl^ trophoblasts, but not in control trophoblasts from *Ifnar^-/-^* mice (**Fig. S13B**). To assess the loss of NPC1 functionality in Ada^cre^/Npc1^fl/fl^ trophoblasts, control and Ada^cre^/Npc1^fl/fl^ trophoblasts were stained with filipin that binds to accumulated cholesterol within lysosomes of NPC1-null cells. Filipin staining in the Ada^cre^/Npc1^fl/fl^ trophoblasts demonstrated increased accumulation indicating decreased NPC1 protein functionality (**Fig. S13C**) (*60*, *61*).

To evaluate the role of NPC1 expression in trophoblasts, E14 Ada^cre^/Npc1^fl/fl^ and control *Ifnar^-/-^* dams were infected with 5×10^6^ TCID_50_ rVSV/EBOV i.v. and maternal, placental, and fetal tissues were harvested on 3 DPI at E17. Maternal serum, liver and spleen titers between *Ifnar^-/-^* and Ada^cre^/Npc1^fl/fl^ mice did not significantly differ (**Fig. 6A-B**), although quantities of infectious virus in liver and serum of the Ada^cre^/Npc1^fl/fl^ mice ranged widely. Titers in the Cre-expressing placentas dropped modestly, but significantly (about 2-fold), providing evidence that JZ trophoblasts contribute to infection levels, but are not the sole placental cell population harboring infection. In fetuses, the decrease was more pronounced with titers decreasing by almost two logs (**Fig. 6C**). A significant difference in the percentage of virus-positive fetuses per dam was found between *Ifnar^-/-^*and Ada^cre^/Npc1^fl/fl^ dams (**Table 2**). Hence, eliminating expression of NPC1 in these trophoblasts modestly reduces the placental viral titers and decreases the frequency of vertical transmission to fetuses, as well as decreases infectious viral titers within fetal liver tissue. In 3 DPI (E17) Ada^cre^/Npc1^fl/fl^ placentas, viral antigen positive cells were assessed by staining for virally expressed GFP. A stark decrease in the presence of this viral antigen was evident in the junctional zone compared to *Ifnar^-/-^* mice (**Fig. 6D**). Focal regions of GFP positivity were present in CK7+ cells within the labyrinth of Ada^cre^/Npc1^fl/fl^ placentas, consistent with specific expression of the Cre recombinase within the junctional zone (**Fig. S13D inset**).

**Fig. 6:**
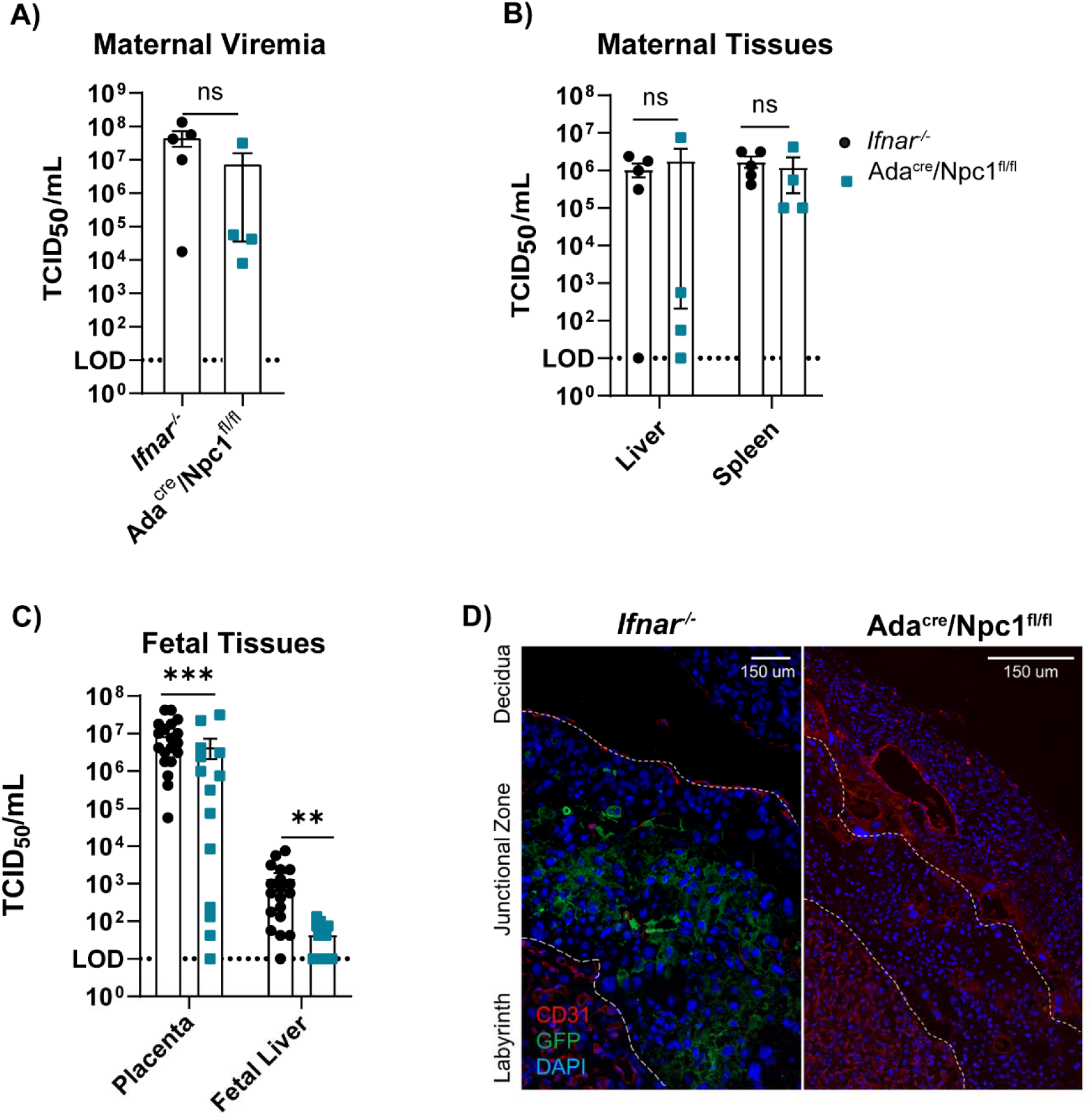
Knock out of NPC1 in junctional zone trophoblasts decreases placental infection and vertical transmission following i.v. administration of rVSV/EBOV. A-C) *lfnar*^-/-^*·* or Ada^cre^/NPC1^fllfl^//fnar^-/-^· dams were infected with 5×10^6^ TCID_50_ of rVSV/EBOV i.v. at E14. **(A)** Maternal sera, **(B)** maternal tissues or **(C)** fetal tissues were harvested at 3 DPI and viral titers quantified by TCID_50_ assays in Vero E6 cells. I*fnar*^-/-^*·*n = 5 dams, Ada^cre^/NPC1^fllff^/Ifnar^-/-^ n = 4 dams.**D)** lmmunostaining of 3 DPI placental sections from /fnar^-/-^·(left) or Ada^cre^/NPC1^fllf^1/Ifnar^-/-^ (right) dam challenged i.v. with rVSV/EBOV. Sections are stained for viral antigen (GFP) and endothelial marker (CD31) and dotted lines demarcate placental regions (20X tilescan). Brightness and contrast uniformly applied across images. ns = not significant, *p≤0.05, **p≤0.01, ***p≤0.001, ****p≤0.0001 as determined by **(A)** unpaired Student’s I-test, **(B-C)** two-way ANOVA w/ Sidak’s multiple comparisons test.

**Table 2:**
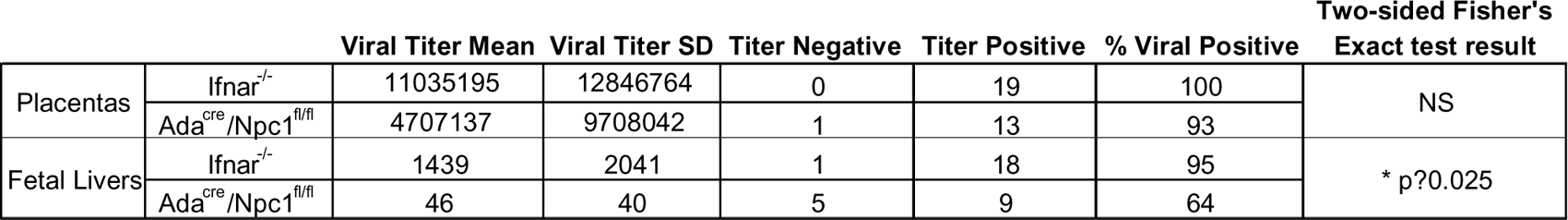
NPC1 expression in junctional zone trophoblasts improves vertical transmission rates. Detection of rVSV/EBOV in placental and fetal livers of Ifnar^-/-^ and Ada^cre^/Npc1^fl/fl^ mice at 3 DPI. Dams were infected with 5×10^6^ TCID_50_ of rVSV/EBOV i.v. on E14. For each tissue, titer mean and standard deviation were calculated. Additionally, tissues were divided by whether they produced infectious viral titer or not. NS = not significant, *p≤0.05 via two-sided Fisher’s Exact test was calculated on these

|  |  | Viral Titer Mean | Viral Titer SD | Titer Negative | Titer Positive | % Viral Positive | Two-sided Fisher's Exact test result |
| --- | --- | --- | --- | --- | --- | --- | --- |
| Placentas | *lfnar<sup>-/-</sup>* | 11035195 | 12846764 | 0 | 19 | 100 | NS |
|  | *Ada<sup>cre</sup>/Npc1<sup>fl/fl</sup>* | 4707137 | 9708042 | 1 | 13 | 93 |  |
| Fetal Livers | *lfnar<sup>-/-</sup>* | 1439 | 2041 | 1 | 18 | 95 | * p?0.025 |
|  | *Ada<sup>cre</sup>/Npc1<sup>fl/fl</sup>* | 46 | 40 | 5 | 9 | 64 |  |

## Discussion

Available data indicate that EBOV infection of pregnant women almost inevitably leads to fetal demise (*21*, *22*). Viral RNA has been reported to be present in ∼96% of tissues evaluated from fetuses from EBOV-positive mothers, suggesting that placental and/or fetal infection leads to loss of the fetus (*22*, *26*). However, little is known about the timing or route of transmission of EBOV from mother to offspring and which cells within placenta and fetus become infected, limiting the understanding of viral pathogenesis during pregnancy. Additionally, it is currently not known if vertical transmission occurs during gestation or during the intrapartum process. Thus, we sought to establish a small animal model to evaluate the vertical transmission of EBOV.

Since experimentation with EBOV required maximum containment facilities, initial studies establishing the model were performed using rVSV/EBOV during timed pregnancies. During late pregnancy, we evaluated several routes of virus administration, finding that i.p., i.v., and i.m. delivery led to maternal and placental infection with delayed but significant fetal infection. These studies provide evidence of *in utero* vertical transmission. Our investigations suggested that i.p. administration likely transmits virus directly from the peritoneum to uterine tissue. As a consequence, placental titers were higher at 1 DPI than titers in maternal visceral organs. Both i.v. and i.m. routes of delivery better modeled a hematogenous route of infection, thereby more accurately mimicking viremic spread of infection in humans.

We observed systemic spread of EBOV in pregnant and non-pregnant females following i.m. infection, with roughly equivalent titers and viral loads irrespective of pregnancy status. Using this route, we show that EBOV can be transmitted *in utero* to the fetus. By 3 DPI when we first evaluated the EBOV-infected mice, maternal viremia was high and was maintained through 5 DPI. At 3DPI, greater than 90% of placentas had detectable virus titers that increased at day 4 of infection. Transmission of EBOV to fetuses was delayed and even by 5 DPI only 10-13% of fetuses had detectable infectious titers, suggesting that placenta tissue provides a partial barrier to fetal transmission. Later time points during EBOV infection should be assessed in the future. While infectious EBOV in fetuses at day 5 of infection was modest, more robust levels of viral RNA within fetal tissues were detected. High levels of RNA present during other *in utero* viral infections are known to elicit immune responses and are associated with pathological outcomes in resulting neonates (*62*). Future experiments should examine alternative viral inoculation doses, alternative administration routes, and longer infection durations.

We found that placental trophoblasts and to a lesser extent placental endothelial cells stained for EBOV antigen. In contrast, placental cells staining for the macrophage marker F4/80 were not EBOV infected. This suggests that placental macrophages do not contribute significantly to virus titers in this organ and are likely not involved in transmission of virus to fetuses. This finding was surprising as macrophages and other myeloid cells such as DCs are well established to be early and sustained targets of EBOV infection in most tissues (*15*, *20*, *63*, *64*). Immunostaining of placentas infected with rVSV/EBOV also demonstrated viral antigen abundantly present in trophoblasts and some endothelial cells. With our low containment virus, we also observed occasional co-staining of viral antigen and IBA1 in infected placentas. This difference may reflect the greater viral antigen abundance in rVSV/EBOV infected placentas.

Highly abundant trophoblasts play many roles in placental function and the development of the fetus, with a primary role to serve as a protective physical and immunological barrier to prevent pathogen transmission from the parent (*34*, *62*, *65*). Since our studies demonstrated that this cell type was frequently infected with virus, we developed a novel mouse strain that eliminated expression of the EBOV receptor, NPC1, in JZ trophoblasts and infected these mice with rVSV/EBOV. At 3 DPI, lower placental and fetal viral titers were evident in these conditional NPC1 knock out mice, as well as lower rates of virus transmission to offspring. Immunostaining of infected placentas of Ada^cre^/Npc1^fl/fl^ mice showed few-to-no viral antigen positive cells within the decidua and junctional zone. However, trophoblasts within the LZ adjacent to the chorionic plate did co-stain for viral antigen, indicating selective deletion of NPC1 in the JZ. These data are consistent with JZ trophoblasts being a critical cell population of EBOV tropism and vertical transmission; however, additional placental cells must also support infection and facilitate EBOV vertical transmission. JZ trophoblasts have also been shown to be abundantly infected by ZIKV during mid-gestation (*66*).

Pro-inflammatory cytokines/chemokines during EBOV infection were elevated in maternal and fetal tissues, consistent with the cytokine storm observed in EBOV-infected patients and experimentally infected animals (*1*, *2*, *67*, *68*). With CXCL10 and CCL2, levels were elevated in maternal serum at earlier times during infection in pregnant dams compare to non-pregnant counterparts, suggesting more rapid initiation of a cytokine levels in the infected pregnant females. RNA of many of these same inflammatory markers were elevated in placental and fetal tissue. This was particularly evident with IL-6, where expression of this cytokine was elevated ∼100-fold in placenta and fetal tissues over the uninfected, pregnant control mice. Elevated levels of IL-6, IL-1, TNF-α and IFN-γ are all associated with negative pregnancy outcomes in human pregnancies, particularly in the context of viral infection (IAV and ZIKV), and parasitic infections (T. gondii) (*45*, *46*, *50*, *69*, *70*). Robust upregulation of the type III interferon IFN-λ in placentas from EBOV-infected pregnancies was also observed compared to naïve placentas. In the context of ZIKV infection, IFN-λ can protect against vertical transmission but may also facilitate negative gestational outcomes (*52*, *53*, *71*, *72*). The consequences of the elevated cytokines within fetal tissues elicited by EBOV need to be more closely examined in future studies.

We utilized *Ifnar^-/-^* mice to achieve robust systemic rVSV/EBOV infections and to enable delivery of authentic EBOV intramuscularly, but this model does not provide insights the role of type I IFNs in control of virus infection or exacerbation of cytokine-induced pathology. Notably, type I IFN signaling presents a dual role: it is implicated in controlling systemic EBOV infection (*73–75*), yet it also acts as a primary mediator of placental damage and fetal loss (*50–52*, *76*). (*37*, *75*, *77*). Fetal type I IFN signaling results in fetal pathology in ZIKV models, and thus we may not be capturing the full scope of fetal pathogenesis in the use of *Ifnar^-/-^* mice (*51*). Future EBOV studies assessing fetal tissues in WT, *Ifnar^+/-^* and *Ifnar^-/-^* dams crossed with males of these different genotypes will provide important insights.

Recent outbreaks of EBOV, BDBV, SARS-CoV-2, ZIKV and hantavirus have caught healthcare systems unprepared, and this may be particularly true for appropriate care of pregnant patients. The most recent large outbreak of BDBV in the Democratic Republic of the Congo and Uganda highlights the critical and ongoing need to understand filovirus pathogenesis in pregnancy (*77*). Viral infections in the context of pregnancy can result in higher rates of pathology for both mother and baby due to the general immunosuppressed state of the mother, allowing for the development of her semi-allogenic fetus (*62*, *78*). At the maternal-fetal interface, the balance of maternal tolerance and anti-microbial defense influences pathogen tropism and the pathogen’s ability to target the fetus. We sought to establish this mouse model to allow further investigation into EBOV pathogenesis in pregnancy. Our data highlights the need for higher quality data collection from human EVD pregnancies and the need for further study of filovirus tropism and kinetics in pregnancy to better understand the pathogenesis in mother and offspring. Our model could be applied to other emerging and high-containment pathogens, as these remain woefully understudied in pregnancy.

## Materials and Methods

### Study Design

These studies were designed to develop a tractable small animal model to investigate EBOV infection. Most of these studies were performed on at late term pregnancies and for low containment studies dams were necropsied on days 1, 2 and 3, whereas high containment studies evaluated infection on days 3, 4 and 5. Sample sizes were determined on the basis of our historical experience and consultation of published literature. All replicates are included in the data presented, and animals of the same genotype were assigned to groups randomly. All BSL2 data presented in main figures were compiled from a minimum of three independent experiments; all BSL4 data are derived from a single study. All ABSL2 experiments were conducted in strict accordance with the Animal Welfare Act and the recommendations in the Guide for the Care and Use of Laboratory Animals of the National Institutes of Health (University of Iowa (UI) Institutional Assurance Number: A3021-01) and in accordance to procedures approved by the University of Iowa Animal Care and Use Committee (protocol #1031280 and #4021280). All infectious work with EBOV was performed in the maximum containment laboratories at the Rocky Mountain Laboratories (RML), Division of Intramural Research, National Institute of Allergy and Infectious Diseases, National Institutes of Health. RML is an institution accredited by the Association for Assessment and Accreditation of Laboratory Animal Care International (AAALAC). All procedures followed standard operating procedures (SOPs) approved by the RML Institutional Biosafety Committee (IBC). The study was approved by the RML Animal Care and Use Committee (ACUC). Procedures were conducted in mice anesthetized by trained personnel under the supervision of veterinary staff. Food and water were available *ad libitum*.

### Mice and timed pregnancies

C57BL/6 *Ifnar^-/-^* mice were a generous gift from Dr. John Harty (University of Iowa, Iowa City, IA, USA). Mice used in BSL4 timed pregnancies were sent to Rocky Mountain Laboratories from the University of Iowa. Harem breeding was used where appropriate. One male per three female mice were placed in a cage overnight. The following morning (16-18 hours later), female mice were vaginally examined for the presence of copulation plugs. The presence of a copulation marked E0 and the female was removed from the cage. Both pregnant and non-female pregnant females were randomly assigned to infection or PBS injection treatment groups.

The B6.FVB-Tg (Ada*-cre)5Xiay/J (Ada-Cre (Strain # 036543)) mice were purchased from The Jackson Laboratory and crossed onto a C57BL/6 *Ifnar^-/-^* backbone in house (Ada^cre^/Ifnar^-/-^). Mice with homozygous floxed *Npc1* alleles (Npc^fl/fl^) were a generous gift from Dr. Andrew Lieberman (University of Michigan) and were also crossed onto an *Ifnar^-/-^* backbone in-house (Npc1^fl/fl^/Ifnar^-/-^) (*79*). Once strains were on *Ifnar^-/-^* backbones, Npc^fl/fl^/*Ifnar^-/-^* mice were crossed with Ada^cre^/*Ifnar^-/-^* mice to create Ada^cre^/ Npc1^fl/fl^/*Ifnar^-/-^* lines. To genotype tissues, we took tail snips and/or ear punches as per IACUC protocols. Genomic DNA was isolated using Promega Wizard Genomic DNA purification kit. Primers utilized are found in **Table S1**

### Viruses

#### BSL2

We utilized rVSV with its native glycoprotein and a GFP reporter (rVSV/G), or with the native glycoprotein replaced with EBOV GP and a GFP reporter (rVSV/EBOV). The propagation of both viruses was performed on mycoplasma-negative Vero E6 and were titered for median tissue culture infectious doses (TCID_50_) assays on Vero E6 cells. Viruses were stored at −80°C until use (*33*).

#### BSL4

EBOV-Mayinga was propagated on mycoplasma negative Vero E6 cells, titered via focus-forming or median tissue culture infectious doses (TCID_50_) assays on Vero E6 cells and stored in liquid nitrogen until use (*80*).

### Viral infection of mice

#### BSL2

For all rVSV stocks used, we performed a dose response curve in non-pregnant *Ifnar^-/-^* females to identify lethal doses of 50% of mice (LD_50_), lethal doses of 75% of mice (LD_75_) and lethal dose for 100% of mice (LD_100_) for each infection route. Survival curves for each infection route were performed for each new stock of virus. Unless otherwise stated, experimental mice were injected with a dose of virus that resulted in an LD_75_ of non-pregnant mice with rVSV/EBOV or rVSV/G. For i.p. administration, an LD_75_ for non-pregnant females was ∼500 TCID_50_. For i.v. and i.m inoculations, the LD_50_ of non-pregnant females was ∼5 x 10^6^ TCID_50_. I.p. inoculations were performed by diluting virus to desired concentration in sterile PBS in a final volume of 100 μl and administrated at the abdominal midline. I.v. doses were diluted in PBS to final volume of 200 μl and delivered retro-orbitally via tuberculin syringe. I.m. injections were given in right rear quadriceps muscle in final volume of 100 μl.

#### BSL4

Mice were infected i.m. with 1×10^4^ FFU on E13 (pregnant) or age matched non-pregnant females. Mice were infected with 100 μl by the i.m. route (50 μl per leg).

### Mouse tissue collection

Mice were euthanized per protocol. Gravid uteri were removed and placed in ice-cold sterile PBS. Using 20 ml of 4°C sterile PBS maternal tissues were perfused via cardiac puncture. Maternal livers and spleens were harvested, weighed, and homogenized in either sterile PBS (viral titer quantification) or TRIzol (RNA solation). Amniotic fluid was removed from each conceptus via tuberculin syringe and pooled per pregnancy. Individual fetoplacental units were isolated from gravid uteri. Fetal membranes and umbilical cords were removed and placentas and fetuses were weighed individually. Tissues were divided for viral titer quantification, RNA isolation, and tissue fixation as appropriate. For fetal tissue viral titrations, umbilical cords were sectioned from both fetal umbilicus and chorionic plates prior to homogenization. Fetal organs were dissected and individually homogenized in PBS.

### Viral tissue titrations

#### BSL2

Tissue homogenates were filtered via 0.45 μm PVDF syringe-tip filters and sterile 1 ml syringes. 10-fold dilutions of homogenate were added to confluent 96 well plates of Vero E6 cells. GFP positivity was scored on 4 DPI. TCID_50_ values were calculated utilizing the Improved Spearman-Karber method (*81*). For some experiments, tissues were weighed in milligrams and weights converted to grams. Where appropriate TCID_50_/ml values were normalized per gram of tissue.

#### BSL4

For determination of EBOV-Mayinga titers in mouse blood and tissue samples, Vero E6 cells were seeded in 48-well plates the day before titration. Tissues were homogenized in 1 mL of DMEM and tissue and blood samples were serial diluted 10-fold. Media was removed from cells and triplicates were inoculated with each dilution. After one hour, DMEM supplemented with 2% FBS, penicillin/streptomycin and L-glutamine was added and incubated at 37°C. Cells were monitored for cytopathic effect (CPE) for 12-14 days and the TCID_50_ was calculated for each sample employing the Reed and Muench method (*82*).

### Viral load and cytokine/chemokine transcript quantification

Tissues were homogenized in TRIzol reagent and total RNA was extracted according to the manufacturer’s instructions. 1 µg of RNA was reverse-transcribed into cDNA using the High-Capacity cDNA Reverse Transcription Kit. Quantitative PCR (qPCR) was then performed using POWER SYBR Green Master Mix following the manufacturer’s protocol. Reactions were run on a QuantStudio 3 Real-Time PCR (Applied Biosystems) system, and cycle threshold (Ct) values were determined using QuantStudio Data Analysis Software. Transcript levels were calculated using the average Ct values from duplicate wells and normalized to the housekeeping gene, mGAPDH, using the 2^−ΔΔCt^ method (*83*). Primer sequences (synthesized by Integrated DNA Technologies, Coralville, IA) are listed in **Table S2**.

### EBOV-Mayinga viral RNA quantification

Levels of EBOV RNA in blood and tissue samples were determined using a probe-based RT-qPCR assay specific to EBOV polymerase (L) as previously described (*84*). RNA from EBOV-Mayinga stock was extracted in the same way as experimental samples and used alongside samples as standards with known TCID_50_ concentrations.

### Tissue staining and immunofluorescence

Direct immunofluorescence was performed on 5 μm sections formalin fixed/paraffin embedded (FFPE) placental tissues attached to glass slides. Tissue sections were deparaffinized by immersion in xylene and rehydrated through graded ethanol series (100%, 95%, 70%, and 50%). Slides were rinsed in ddH_2_0. Antigen retrieval was performed in 10 mM citrate buffer (pH 6.0) with 1 cycle at 60°C for 1 minute followed by 2 cycles at 90°C for 5 minutes each, with a 5-minute cooling period between. Tissues were permeabilized using 0.5% Triton X-100 in 1% TBS. Slides were washed and then blocked in immunofluorescence buffer containing 5% donkey or goat serum, 5% BSA, and 0.1% fish skin gelatin. After blocking, primary antibodies diluted in blocking buffer were added and slides were incubated in a humidified chamber. Antibody dilutions and incubation conditions used is in **Table S3**. Slides were washed thrice in with 1X PBS containing 0.015% Tween 20. When needed, sections were incubated at RT for 1 hour with species-specific secondary antibodies diluted in blocking buffer (25 μl) diluted in blocking buffer. Slides were washed thrice and mounted with Thermo Prolong Diamond Antifade mounting medium with DAPI. Image acquisition was performed using a Zeiss confocal microscope Zen Blue acquisition software. In certain instances, entire images were cropped or underwent color adjustment using ImageJ.

### Co-localization analysis

Co-localization of viral antigen and tissue markers was analyzed using the *Co-localization Highlighter* plugin from the MBF ImageJ plugin collection (https://imagej.net/ij/plugins/mbf/index.html). Following channel separation, threshold values for the green (viral antigen) and red (tissue markers) channels were established from control images and applied uniformly across all datasets. Overlapping signals were identified with the Co-localization Highlighter, where colocalized pixels appeared in white. The resulting composite was separated into individual channels, with the colocalization signal isolated in the blue channel. Regions of interest (ROIs) were defined, and the percentage area of colocalization was quantified from the blue channel.

### Serum cytokine quantification

Cytokines and chemokines in serum from experimental animals were assessed in a bead-based ELISA flow cytometry multiplex assay (BioLegend, Cat #: 70622). Serum harvested from EBOV Mayinga infected animals was gamma-irradiated in accordance with RML biosafety protocols. Assay and analysis were performed following manufacturer’s instructions utilizing a Beckman CytoFLEX flow cytometer.

### Trophoblast tissue enzymatic digestion and isolation

Trophoblasts were enzymatically dissociated from either *Ifnar^-/-^*or Ada^cre^/Npc1^fl/fl^/*Ifnar^-/-^* placentas using the protocol described previously (*59*). Briefly, placentas were harvested from 1-2 E15-18 *Ifnar^-/-^* or Ada^cre^/Npc1^fl/fl^/*Ifnar^-/-^* pregnancies. Uterine and fetal membrane tissues were dissected away from the placenta, and placentas minced in ice cold digestion buffer (final concentrations of DNAse I 20 u/ml and 125 digestion units/ml collagenase suspended in 100 ml of wash buffer (Medium 199, 0.02M HEPES, 0.01M sodium bicarbonate, 100ug/ml penicillin/streptomycin)). Tissue was incubated in 37°C water bath for ∼1 hour. Samples were pipetted up and down 5-10 times with a 10 ml pipet every 15 minutes. Samples were strained through a 100 µM strainer into a fresh tube and centrifuged at 500 x g for 5 minutes (m). Supernatant was removed and cells were washed with 10 ml of wash buffer and centrifuged again at 500 x g for 5 minutes. Cells were resuspended and loaded onto a Percoll solution (9.6 ml Percoll, 13.4 ml of wash buffer, and 1.1 ml of 10x Medium 199). Percoll solution with cells was inverted twice and centrifuged at 30,000 x g for 40 minutes at 4°C. Trophoblasts were harvested from the middle layer of the Percoll gradient and washed once with wash buffer. Trophoblasts used for evaluation of cre expression were placed in TRIzol for RNA extraction. Cells utilized for filipin staining were counted and then were plated in RPMI + 10% FBS + 1% P/S and kept in a 37°C CO_2_ incubator with water bath and 5% CO_2_ overnight.

### Intracellular NPC1 staining and quantification

Trophoblasts were washed once with 1x PBS and fixed with 4% PFA at RT for 15 minutes. Cells were washed twice with 1x PBS. Wheat germ agglutinin (WGA) conjugated to Alexa fluor 594 was applied to cells diluted 1:5000 in PBS and was incubated at RT for one hour in the dark. Cells were washed twice again with 1x PBS. Then cells were permeabilized with 0.25% TBS/TritonX-100 for 10 minutes RT. Cells were washed twice with 1x PBS. Filipin solution was prepared fresh each day with 1ug of filipin being diluted into 40 μl of DMSO and added to cells in 1 ml of PBS. After a 10-minute incubation at RT cells were imaged on an inverted fluorescent Nikon Eclipse Ti series microscope at 20x magnification.

### Statistical analysis

Data analysis and figure production were conducted utilizing GraphPad Prism (v. 10.5.0). Where applicable, nonlinear regression and false discovery rate outlier analysis and outlier removal were performed on each data set prior to statistical analyses (*85*). For viral titers or viral loads quantified via qPCR statistical analyses were performed on log transformed data and depicted on a log_10_ scale. Data on graphs are depicted as mean with error bars denoting the S.E.M. Statistical analyses run for each experiment can be found in figure legends. Significance was considered with a p-value of ≤0.05 and degree of significance was depicted with various asterisks as defined in figure legends. All data represented are from at least 2 independent experiments unless otherwise noted.

## Acknowledgments & Funding

This study was supported by NIH NIAID UH2AI186375 (WJM). HVE was supported throughout this work via NIH NIAID F30AI174686, University of Iowa MSTP 5T32 GM139776-03 and University of Iowa Immunology Graduate Program in Immunology T32 AI0077485. Support was also provided by NIH GMR35154959 (MLS). This research was supported in part by the Intramural Research Program of the National Institutes of Health (NIH) (AI001254 to AM).

The contributions of the NIH authors were made as part of their official duties as NIH federal employees, are in compliance with agency policy requirements, and are considered Works of the United States Government. However, the findings and conclusions presented in this paper are those of the author and do not necessarily reflect the views of the NIH or the U.S. Department of Health and Human Services.

The authors would like to acknowledge use of the University of Iowa Central Microscopy Research Facility, a core resource supported by the University of Iowa Vice President for Research, and the Carver College of Medicine. We acknowledge the personnel and instrumentation in the Neural Circuits and Behavior Core in the Iowa Neuroscience Institute, supported in part by the Roy J. Carver Charitable Trust and UI Carver College of Medicine. We also would like to thank the animal care staff of University of Iowa and the Rocky Mountain Veterinary Branch for their support in this study.

## Author contributions

Conceptualization – H.V.E., M.S., A.M., W.J.M.

Data curation – H.V.E., W.J.M.

Formal analysis – H.V.E., C.W.H.

Funding acquisition – H.V.E., A.M., K.M., W.J.M.

Investigation – experimentation H.V.E., E.K., E.E., C.W.H., B.J.S., P.R., M.F., M.L.

Methodology -- H.V.E., C.W.H., B.J.S., K.N.M., A.M., M.L.S., W.J.M.

Project Administration – H.V.E., C.W.H., A.M., W.J.M.

Resources – H.V.E., K.N.M., A.M., W.J.M.

Supervision – H.V.E., K.N.M., A.M., W.J.M.

Visualization – H.V.E., K.N.M., W.J.M.

Writing – original draft – H.V.E., W.J.M.

Writing – reviewing and editing – H.V.E., K.N.M., P.R., C.W.H., K.N.M., M.K.S., A.M., W.J.M.

## References

1. S. T. Jacob, I. Crozier, W. A. Fischer, A. Hewlett, C. S. Kraft, M. A. de La Vega, M. J. Soka, V. Wahl, A. Griffiths, L. Bollinger, J. H. Kuhn, Ebola Virus Disease (Springer US, 2020; 10.1038/s41572-020-0147-3) vol. 6.

2. H. Feldmann, T. W. Geisbert, Ebola haemorrhagic fever. The Lancet 377, 849–862 (2011).

3. V. Vine, D. P. Scott, H. Feldmann, “Ebolavirus: An Overview of Molecular and Clinical Pathogenesis” in Methods in Molecular Biology (Humana Press Inc., 2017; http://link.springer.com/10.1007/978-1-4939-7116-9_3) vol. 1628, pp. 39–50.

4. L. Baseler, D. S. Chertow, K. M. Johnson, H. Feldmann, D. M. Morens, The Pathogenesis of Ebola Virus Disease. Annu. Rev. Pathol. Mech. Dis. 2017 12, 387–418 (2016).

5. WHO, Ebola virus disease Fact Sheet. https://www.who.int/news-room/fact-sheets/detail/ebola-virus-disease.

6. E. O. Musa, E. Adedire, O. Adeoye, P. Adewuyi, N. Waziri, P. Nguku, M. Nanjuya, B. Adebayo, A. Fatiregun, B. Enya, C. Ohuabunwo, K. Sabitu, F. Shuaib, A. Okoh, O. Oguntimehin, N. Onyekwere, A. Nasidi, A. Olayinka, Epidemiological profile of the Ebola virus disease outbreak in Nigeria, July-September 2014. Pan African Medical Journal 21, 1–6 (2015).

7. N. G. Onyeneho, N. I. Aronu, I. Igwe, J. Okeibunor, T. Diarra, J. N. Anoko, M. H. Djingarey, Z. Yoti, D. Chamla, A. S. Gueye, The Impact of the Ebola Virus Disease Epidemic among Women in the Provinces of North Kivu and Ituri in the Democratic Republic of the Congo. J. Immunol. Sci. Suppl 3, 11 (2023).

8. C. Menéndez, A. Lucas, K. Munguambe, A. Langer, Ebola crisis: The unequal impact on women and children’s health. Lancet Glob. Health 3, e130 (2015).

9. P. T. Richards, A. M. Fleck, R. Patel, M. Fakhimi, D. Bohan, K. Geoghegan-Barek, A. N. Honko, A. E. Stolte, C. B. Plescia, C. O. Messingham, S. J. Connell, T. P. Crowe, F. A. Gourronc, R. Carrion, A. Griffiths, D. K. Meyerholz, A. J. Klingelhutz, R. A. Davey, K. N. Messingham, W. Maury, Ebola virus’ hidden target: virus transmission to and infection of skin. J. Virol., doi: 10.1128/jvi.01300-25 (2025).

10. K. N. Messingham, P. T. Richards, A. Fleck, R. A. Patel, M. Djurkovic, J. Elliff, S. Connell, T. P. Crowe, J. Munoz Gonzalez, F. Gourronc, J. A. Dillard, R. A. Davey, A. Klingelhutz, O. Shtanko, W. Maury, Multiple cell types support productive infection and dynamic translocation of infectious Ebola virus to the surface of human skin. Sci. Adv. 11 (2025).

11. M. Côté, J. Misasi, T. Ren, A. Bruchez, K. Lee, C. M. Filone, L. Hensley, Q. Li, D. Ory, K. Chandran, J. Cunningham, Small molecule inhibitors reveal Niemann-Pick C1 is essential for Ebola virus infection. Nature 477, 344–348 (2011).

12. E. H. Miller, G. Obernosterer, M. Raaben, A. S. Herbert, M. S. Deffieu, A. Krishnan, E. Ndungo, R. G. Sandesara, J. E. Carette, A. I. Kuehne, G. Ruthel, S. R. Pfeffer, J. M. Dye, S. P. Whelan, T. R. Brummelkamp, K. Chandran, Ebola virus entry requires the host-programmed recognition of an intracellular receptor. EMBO Journal 31, 1947–1960 (2012).

13. B. M. Connolly, K. E. Steele, K. J. Davis, T. W. Geisbert, W. M. Kell, N. K. Jaax, P. B. Jahrling, Pathogenesis of Experimental Ebola Virus Infection in Guinea Pigs. J. Infect. Dis. 179, S203–S217 (1999).

14. M. Bray, K. Davis, T. Geisbert, C. Schmaljohn, J. Huggins, A Mouse Model for Evaluation of Prophylaxis and Therapy of Ebola Hemorrhagic Fever. J. Infect. Dis. 178, 651–661 (1998).

15. T. R. Gibb, M. Bray, T. W. Geisbert, K. E. Steele, W. M. Kell, K. J. Davis, N. K. Jaax, Pathogenesis of Experimental Ebola Zaire Virus Infection in BALB / c Mice. J. Comp. Pathol. 125, 233–242 (2001).

16. T. W. Geisbert, L. E. Hensley, T. Larsen, H. A. Young, D. S. Reed, J. B. Geisbert, D. P. Scott, E. Kagan, P. B. Jahrling, K. J. Davis, Pathogenesis of Ebola Hemorrhagic Fever in Cynomolgus Macaques. Am. J. Pathol. 163, 2347–2370 (2003).

17. C. M. Bosio, M. J. Aman, C. Grogan, R. Hogan, G. Ruthel, D. Negley, M. Mohamadzadeh, S. Bavari, A. Schmaljohn, Ebola and Marburg Viruses Replicate in Monocyte Derived Dendritic Cells without Inducing the Production of Cytokines and Full Maturation. J. Infect. Dis. 188, 1630–1638 (2003).

18. N. M. Lubaki, P. Ilinykh, C. Pietzsch, B. Tigabu, A. N. Freiberg, R. A. Koup, A. Bukreyev, The Lack of Maturation of Ebola Virus-Infected Dendritic Cells Results from the Cooperative Effect of at Least Two Viral Domains. J. Virol. 87, 7471–7485 (2013).

19. B. Yen, L. C. F. Mulder, O. Martinez, C. F. Basler, Molecular Basis for Ebolavirus VP35 Suppression of Human Dendritic Cell Maturation. J. Virol. 88, 12500–12510 (2014).

20. S. B. Bradfute, K. L. Warfield, M. Bray, Mouse Models for Filovirus Infections. Viruses 4, 1477–1508 (2012).

21. M. E. Foeller, C. Carvalho Ribeiro do Valle, T. M. Foeller, O. T. Oladapo, E. Roos, A. E. Thorson, C. Carvalho, T. M. Foeller, O. T. Oladapo, E. Roos, A. E. Thorson, Review Pregnancy and breastfeeding in the context of Ebola : a systematic review. Lancet Infect. Dis. 20, 149–158 (2020).

22. N. D. Kayem, C. Benson, C. Y. L. Aye, S. Barker, M. Tome, S. Kennedy, P. Ariana, P. Horby, Ebola virus disease in pregnancy: a systematic review and meta-analysis. Trans. R. Soc. Trop. Med. Hyg., 509–522 (2021).

23. K. Appiah-Sakyi, M. Mohan, J. C. Konje, Ebola infection in pregnancy, an ongoing challenge for both the global health expert and the pregnant woman—A review. European Journal of Obstetrics & Gynecology and Reproductive Biology 258, 111–117 (2021).

24. N. S. Olgun, Viral Infections in Pregnancy: A Focus on Ebola Virus. Curr. Pharm. Des. 24, 993–998 (2018).

25. L. M. Bebell, T. Oduyebo, L. E. Riley, Ebola virus disease and pregnancy: A review of the current knowledge of Ebola virus pathogenesis, maternal, and neonatal outcomes. Birth Defects Res. 109, 353–362 (2017).

26. A. Muehlenbachs, O. de la Rosa Vázquez, D. G. Bausch, I. J. Schafer, C. D. Paddock, J. P. Nyakio, P. Lame, E. Bergeron, A. M. McCollum, C. S. Goldsmith, B. C. Bollweg, M. A. Prieto, R. S. Lushima, B. K. Ilunga, S. T. Nichol, W. J. Shieh, U. Ströher, P. E. Rollin, S. R. Zaki, Ebola Virus Disease in Pregnancy: Clinical, Histopathologic, and Immunohistochemical Findings. J. Infect. Dis. 215, 64–69 (2017).

27. S. A. Riesle-Sbarbaro, G. Wibbelt, A. Düx, V. Kouakou, M. Bokelmann, K. Hansen-Kant, N. Kirchoff, M. Laue, N. Kromarek, A. Lander, U. Vogel, A. Wahlbrink, D. M. Wozniak, D. P. Scott, J. B. Prescott, L. Schaade, E. Couacy-Hymann, A. Kurth, Selective replication and vertical transmission of Ebola virus in experimentally infected Angolan free-tailed bats. Nat. Commun. 15 (2024).

28. K. Chandran, N. J. Sullivan, U. Felbor, S. P. Whelan, J. M. Cunningham, Virology: Endosomal proteolysis of the ebola virus glycoprotein is necessary for infection. Science (1979). 308, 1643–1645 (2005).

29. B. A. Rhein, L. S. Powers, K. Rogers, M. Anantpadma, B. K. Singh, Y. Sakurai, T. Bair, C. Miller-Hunt, P. Sinn, R. A. Davey, M. M. Monick, W. Maury, Interferon-γ Inhibits Ebola Virus Infection. PLoS Pathog. 11, 1–28 (2015).

30. B. Brunton, K. Rogers, E. K. Phillips, R. B. Brouillette, R. Bouls, N. S. Butler, W. Maury, TIM-1 serves as a receptor for Ebola virus in vivo, enhancing viremia and pathogenesis. PLoS Negl. Trop. Dis. 13, e0006983 (2019).

31. K. J. Rogers, B. Brunton, L. Mallinger, D. Bohan, K. M. Sevcik, J. Chen, N. Ruggio, W. Maury, IL-4/IL-13 polarization of macrophages enhances Ebola virus glycoprotein-dependent infection. PLoS Negl. Trop. Dis. 13, e0007819 (2019).

32. K. J. Rogers, O. Shtanko, R. Vijay, L. N. Mallinger, C. J. Joyner, M. R. Galinski, N. S. Butler, W. Maury, Acute Plasmodium Infection Promotes Interferon-Gamma-Dependent Resistance to Ebola Virus Infection. Cell Rep. 30, 4041–4051.e4 (2020).

33. P. T. Richards, J. A. A. Briseño, B. A. Brunton, W. Maury, “In Vivo Investigation of Filovirus Glycoprotein-Mediated Infection in a BSL2 Setting” (2025; https://link.springer.com/10.1007/978-1-0716-4256-6_13), pp. 183–198.

34. S. A. Elmore, R. Z. Cochran, B. Bolon, B. Lubeck, B. Mahler, D. Sabio, J. M. Ward, Histology Atlas of the Developing Mouse Placenta. Toxicol. Pathol. 50, 60–117 (2022).

35. Q. H. Li, K. Kim, S. Shresta, Mouse models of Zika virus transplacental transmission. Antiviral Res. 210, 105500 (2023).

36. A. L. Fowden, A. N. Sferruzzi-Perri, P. M. Coan, M. Constancia, G. J. Burton, Placental efficiency and adaptation: endocrine regulation. J. Physiol. 587, 3459 (2009).

37. J. R. Spengler, K. J. Lavender, C. Martellaro, A. Carmody, A. Kurth, J. G. Keck, G. Saturday, D. P. Scott, S. T. Nichol, K. J. Hasenkrug, C. F. Spiropoulou, H. Feldmann, J. Prescott, Ebola Virus Replication and Disease Without Immunopathology in Mice Expressing Transgenes to Support Human Myeloid and Lymphoid Cell Engraftment. Journal of Infectious Diseases 214, S308–S318 (2016).

38. M. Hemberger, C. W. Hanna, W. Dean, Mechanisms of early placental development in mouse and humans. Nat. Rev. Genet. 21 (2020).

39. A. Malassiné, J. L. Frendo, D. Evain-Brion, A comparison of placental development and endocrine functions between the human and mouse model. Hum. Reprod. Update 9, 531–539 (2003).

40. D. G. Simmons, A. L. Fortier, J. C. Cross, Diverse subtypes and developmental origins of trophoblast giant cells in the mouse placenta. Dev. Biol. 304, 567–578 (2007).

41. S. J. Tunster, E. D. Watson, A. L. Fowden, G. J. Burton, Placental glycogen stores and fetal growth: insights from genetic mouse models. Reproduction 159, R213–R235 (2020).

42. G. Liang, C. Zhou, X. Jiang, H. Wang, J. J. Han, F. Liu, Article De novo generation of macrophage from placenta- derived hemogenic endothelium ll Article De novo generation of macrophage from placenta-derived hemogenic endothelium. Dev. Cell 56, 2121–2133.e6 (2021).

43. X. Chen, A. T. Tang, J. Tober, J. Yang, N. A. Leu, S. Sterling, M. Chen, Y. Yang, P. Mericko-Ishizuka, N. A. Speck, M. L. Kahn, Mouse placenta fetal macrophages arise from endothelial cells outside the placenta. Dev. Cell 57, 2652–2660.e3 (2022).

44. T. W. Geisbert, H. A. Young, P. B. Jahrling, K. J. Davis, T. Larsen, E. Kagan, L. E. Hensley, Pathogenesis of Ebola Hemorrhagic Fever in Primate Models: Evidence that Hemorrhage Is Not a Direct Effect of Virus-Induced Cytolysis of Endothelial Cells. American Journal of Pathology 163, 2371–2382 (2003).

45. A. Senegas, O. Villard, A. Neuville, L. Marcellin, A. W. Pfaff, T. Steinmetz, M. Mousli, J. P. Klein, E. Candolfi, Toxoplasma gondii-induced foetal resorption in mice involves interferon-gamma-induced apoptosis and spiral artery dilation at the maternofoetal interface. Int. J. Parasitol. 39, 481–487 (2009).

46. T. Srivastava, T. Joshi, D. P. Heruth, M. H. Rezaiekhaligh, R. E. Garola, J. Zhou, V. C. Boinpelly, M. F. Ali, U. S. Alon, M. Sharma, G. B. Vanden Heuvel, P. Mahajan, L. Priya, Y. Jiang, E. T. McCarthy, V. J. Savin, R. Sharma, M. Sharma, A mouse model of prenatal exposure to Interleukin-6 to study the developmental origin of health and disease. Sci. Rep. 11, 13260 (2021).

47. N. Elemam, I. Talaat, A. Maghazachi, CXCL10 Chemokine: A Critical Player in RNA and DNA Viral Infections. Viruses 14, 2445 (2022).

48. B. Wolszczak-Biedrzycka, B. Cieślikiewicz, F. Studniarz, Ł. Dąbrowski, M. Fąs, K. Matyszkiewicz–Suchodolska, M. Harasimowicz, J. Dorf, Chemokines as potential biomarkers for predicting the course of COVID-19 – a review of the literature. Front. Immunol. 16, 1–12 (2025).

49. J. Korbecki, M. Bosiacki, I. Szatkowska, P. Kupnicka, D. Chlubek, I. Baranowska-Bosiacka, The Clinical Significance and Involvement in Molecular Cancer Processes of Chemokine CXCL1 in Selected Tumors. Int. J. Mol. Sci. 25, 1–28 (2024).

50. A. T. Harding, M. A. Goff, H. M. Froggatt, J. K. Lim, N. S. Heaton, GPER1 is required to protect fetal health from maternal inflammation. Science (1979). 371, 271–276 (2021).

51. L. J. Yockey, K. A. Jurado, N. Arora, A. Millet, T. Rakib, K. M. Milano, A. K. Hastings, E. Fikrig, Y. Kong, T. L. Horvath, S. Weatherbee, H. J. Kliman, C. B. Coyne, A. Iwasaki, Type I interferons instigate fetal demise after Zika virus infection. Sci. Immunol. 3 (2018).

52. R. L. Casazza, D. T. Philip, H. M. Lazear, Interferon Lambda Signals in Maternal Tissues to Exert Protective and Pathogenic Effects in a Gestational Stage-Dependent Manner. mBio 13 (2022).

53. M. R. Dedloff, H. M. Lazear, Antiviral and Immunomodulatory Effects of Interferon Lambda at the Maternal-Fetal Interface. Annu. Rev. Virol. 11, 363–379 (2024).

54. N. Sharma, C. Kubaczka, S. Kaiser, D. Nettersheim, S. S. Mughal, S. Riesenberg, M. Hölzel, E. Winterhager, H. Schorle, Tpbpa-Cre-mediated deletion of TFAP2C leads to deregulation of Cdkn1a, Akt1 and the ERK pathway, causing placental growth arrest. Development (Cambridge) 143, 787–798 (2016).

55. D. Hu, J. C. Cross, Ablation of Tpbpa-positive trophoblast precursors leads to defects in maternal spiral artery remodeling in the mouse placenta. Dev. Biol. 358, 231–239 (2011).

56. C. C. Zhou, J. Chang, T. Mi, S. Abbasi, D. Gu, L. Huang, W. Z. Zhang, R. E. Kellems, R. J. Schwartz, Y. Xia, Targeted expression of Cre recombinase provokes placental-specific DNA recombination in transgenic mice. PLoS One 7 (2012).

57. M. J. Elrick, C. D. Pacheco, T. Yu, N. Dadgar, V. G. Shakkottai, C. Ware, H. L. Paulson, A. P. Lieberman, Conditional Niemann-Pick C mice demonstrate cell autonomous Purkinje cell neurodegeneration. Hum. Mol. Genet. 19, 837–847 (2010).

58. R. Spiegel, A. Raas-Rothschild, O. Reish, M. Regev, V. Meiner, R. Bargal, V. Sury, K. Meir, M. Nadjari, G. Hermann, T. C. Iancu, S. A. Shalev, M. Zeigler, The clinical spectrum of fetal Niemann-Pick type C. Am. J. Med. Genet. A 149, 446–450 (2009).

59. K. A. Pennington, J. M. Schlitt, L. C. Schulz, Isolation of primary mouse trophoblast cells and trophoblast invasion assay. Journal of Visualized Experiments, 1–6 (2012).

60. W. Drabikowski, E. Łagwińska, M. G. Sarzała, Filipin as a fluorescent probe for the location of cholesterol in the membranes of fragmented sarcoplasmic reticulum. Biochimica et Biophysica Acta (BBA) - Biomembranes 291, 61–70 (1973).

61. E. J. Blanchette-Mackie, Intracellular cholesterol trafficking: role of the NPC1 protein. Biochim. Biophys. Acta 1486, 171–183 (2000).

62. A. Espino, H. El Costa, J. Tabiasco, R. Al-Daccak, N. Jabrane-Ferrat, Innate Immune Response to Viral Infections at the Maternal-Fetal Interface in Human Pregnancy. Front. Med. (Lausanne). 8, 1–9 (2021).

63. M. Bray, T. W. Geisbert, Ebola virus: The role of macrophages and dendritic cells in the pathogenesis of Ebola hemorrhagic fever. Int. J. Biochem. Cell Biol. 37, 1560–1566 (2005).

64. T. W. Geisbert, P. B. Jahrling, M. A. Hanes, P. M. Zack, Association of Ebola-related Reston Virus Particles and Antigen with Tissue Lesions of Monkeys Imported to the United States. J. Comp. Pathol. 106, 137–152 (1992).

65. E. F. Cornish, I. Filipovic, F. Åsenius, D. J. Williams, T. McDonnell, Innate Immune Responses to Acute Viral Infection During Pregnancy. Front. Immunol. 11 (2020).

66. Y. Alippe, L. Wang, R. Coskun, S. P. Muraro, F. R. Zhao, M. Elam-Noll, J. M. White, D. M. Vota, V. C. Hauk, J. I. Gordon, S. A. Handley, M. S. Diamond, Fetal MAVS and type I IFN signaling pathways control ZIKV infection in the placenta and maternal decidua. Journal of Experimental Medicine 221 (2024).

67. L. Banadyga, A. Marzi, J. Lu, J. M. Gullett, T.-D. Kanneganti, Filoviruses: Innate Immunity, Inflammatory Cell Death, and Cytokines. doi: 10.3390/pathogens11121400 (2022).

68. V. Siragam, G. Wong, X.-G. Qiu, Animal Models for Filovirus Infections. Zool. Res. 39, 15–24 (2018).

69. S. P. Murphy, C. Tayade, A. A. Ashkar, K. Hatta, J. Zhang, B. A. Croy, Interferon Gamma in Successful Pregnancies. Biol. Reprod. 80, 848–859 (2009).

70. J. R. Prins, N. Gomez-Lopez, S. A. Robertson, Interleukin-6 in pregnancy and gestational disorders. J. Reprod. Immunol. 95, 1–14 (2012).

71. A. I. Wells, C. B. Coyne, Type III Interferons in Antiviral Defenses at Barrier Surfaces. Trends Immunol. 39, 848–858 (2018).

72. N. J. Bayer, A. Lennemann, Y. Ouyang, J. C. Bramley, S. Morosky, E. T. Marques Jr., S. Cherry, Y. Sadovsky, Coyne. C. B., Type III Interferons Produced by Human Placental Trophoblasts Confer Protection Against Zika Virus Infection. Cell Host Microbe 19, 1–17 (2016).

73. M. Bray, The role of the type I interferon response in the resistance of mice of filovirus infection. Journal of General Virology 82, 1365–1373 (2001).

74. J. R. Spengler, S. R. Welch, J. M. Ritter, J. R. Harmon, J. D. Coleman-McCray, S. C. Genzer, J. N. Seixas, F. E. M. Scholte, K. A. Davies, S. B. Bradfute, J. M. Montgomery, C. F. Spiropoulou, Mouse models of Ebola virus tolerance and lethality: characterization of CD-1 mice infected with wild-type, guinea pig-adapted, or mouse-adapted virus. Antiviral Res. 210, 105496 (2023).

75. M. S. Lever, T. J. Piercy, J. A. Steward, L. Eastaugh, S. J. Smither, C. Taylor, F. J. Salguero, R. J. Phillpotts, Lethality and pathogenesis of airborne infection with filoviruses in A129 α/β -/- interferon receptor-deficient mice. J. Med. Microbiol. 61, 8–15 (2012).

76. L. J. Yockey, A. Iwasaki, Interferons and Proinflammatory Cytokines in Pregnancy and Fetal Development. Immunity 49, 397–412 (2018).

77. World Health Organization, Disease Outbreak News; Bundibugyo Virus Disease, Democratic Republic of the Congo and Uganda (2026).

78. C. J. Megli, C. B. Coyne, Infections at the maternal–fetal interface: an overview of pathogenesis and defence. Nat. Rev. Microbiol. 20, 67–82 (2022).

79. M. J. Elrick, C. D. Pacheco, T. Yu, N. Dadgar, V. G. Shakkottai, C. Ware, H. L. Paulson, A. P. Lieberman, Conditional Niemann-Pick C mice demonstrate cell autonomous Purkinje cell neurodegeneration. Hum. Mol. Genet. 19, 837–847 (2010).

80. A. Marzi, F. Feldmann, P. W. Hanley, D. P. Scott, S. Günther, H. Feldmann, Delayed disease progression in cynomolgus macaques infected with ebola virus makona strain. Emerg. Infect. Dis. 21, 1777–1783 (2015).

81. C. Lei, J. Yang, J. Hu, X. Sun, On the Calculation of TCID50 for Quantitation of Virus Infectivity. Virol. Sin. 36, 141 (2020).

82. L. J. Reed, H. Muench, A Simple Method of Estimating Fifty Percent Endpoints. Am. J. Epidemiol. 27, 493–497 (1938).

83. K. J. Livak, T. D. Schmittgen, Analysis of Relative Gene Expression Data Using Real-Time Quantitative PCR and the 2−ΔΔCT Method. Methods 25, 402–408 (2001).

84. A. Domi, F. Feldmann, R. Basu, N. McCurley, K. Shifflett, J. Emanuel, M. S. Hellerstein, F. Guirakhoo, C. Orlandi, R. Flinko, G. K. Lewis, P. W. Hanley, H. Feldmann, H. L. Robinson, A. Marzi, A Single Dose of Modified Vaccinia Ankara expressing Ebola Virus Like Particles Protects Nonhuman Primates from Lethal Ebola Virus Challenge. Sci. Rep. 8, 1–9 (2018).

85. H. J. Motulsky, R. E. Brown, Detecting outliers when fitting data with nonlinear regression – a new method based on robust nonlinear regression and the false discovery rate. BMC Bioinformatics 7, 123 (2006).

