## Supplemental figures and tables for "Developing and Characterizing a Murine Model of *In Utero* Transmission of Ebola Virus"

**List of Supplemental Materials**

Fig. S1: Characterizing rVSV/EBOV kinetics in pregnant and non-pregnant *Ifnar^-/-^* mice infected intraperitoneally

Fig. S2: EBOV GP, but not VSV G, facilitates placental infection and vertical transmission to fetuses at 3 DPI

Fig. S3: Impact of i.p. rVSV/EBOV infection on maternal vs fetal tissues and outcomes

Fig. S4: rVSV/EBOV infection i.p. on E5 results in infected concepti tissue and reduced conceptus weights at 3 DPI

Fig. S5: Intravenous rVSV/EBOV administered at E14 does not influence fetal outcomes

Fig. S6: Subcutaneous (sub.q.) and intravaginal (i.vag.) routes of rVSV/EBOV inoculation do not result in robust systemic spread of virus.

Fig. S7: Intramuscular (i.m.) infection of pregnant Ifnar^-/-^ dams with rVSV/EBOV results in maternal, placental, and fetal infection with mild impacts on fetal outcomes

Fig. S8: EBOV titers or viral loads are not altered by pregnancy

Fig. S9: rVSV/EBOV has similar distribution to EBOV within *Ifnar^-/-^* placental samples

Fig. S10: Maternal serum has increasing concentrations of pro-inflammatory cytokines and chemokines upon i.m. inoculation of EBOV

Fig. S11: Elevated pro-inflammatory cytokines and chemokines in sera from rVSV/EBOV infected dams.

Fig. S12: Pro-inflammatory cytokines and chemokine transcripts are elevated in EBOV infected placenta and fetal tissues

Fig. S13: Characterization of Ada^cre^/Npc1^fl/fl^/*Ifnar^-/-^* mouse strain.

Table S1: Murine genotyping primers

Table S2: qPCR primer sequences

Table S3: Antibodies, optimal dilutions, and incubation conditions for immunofluorescence

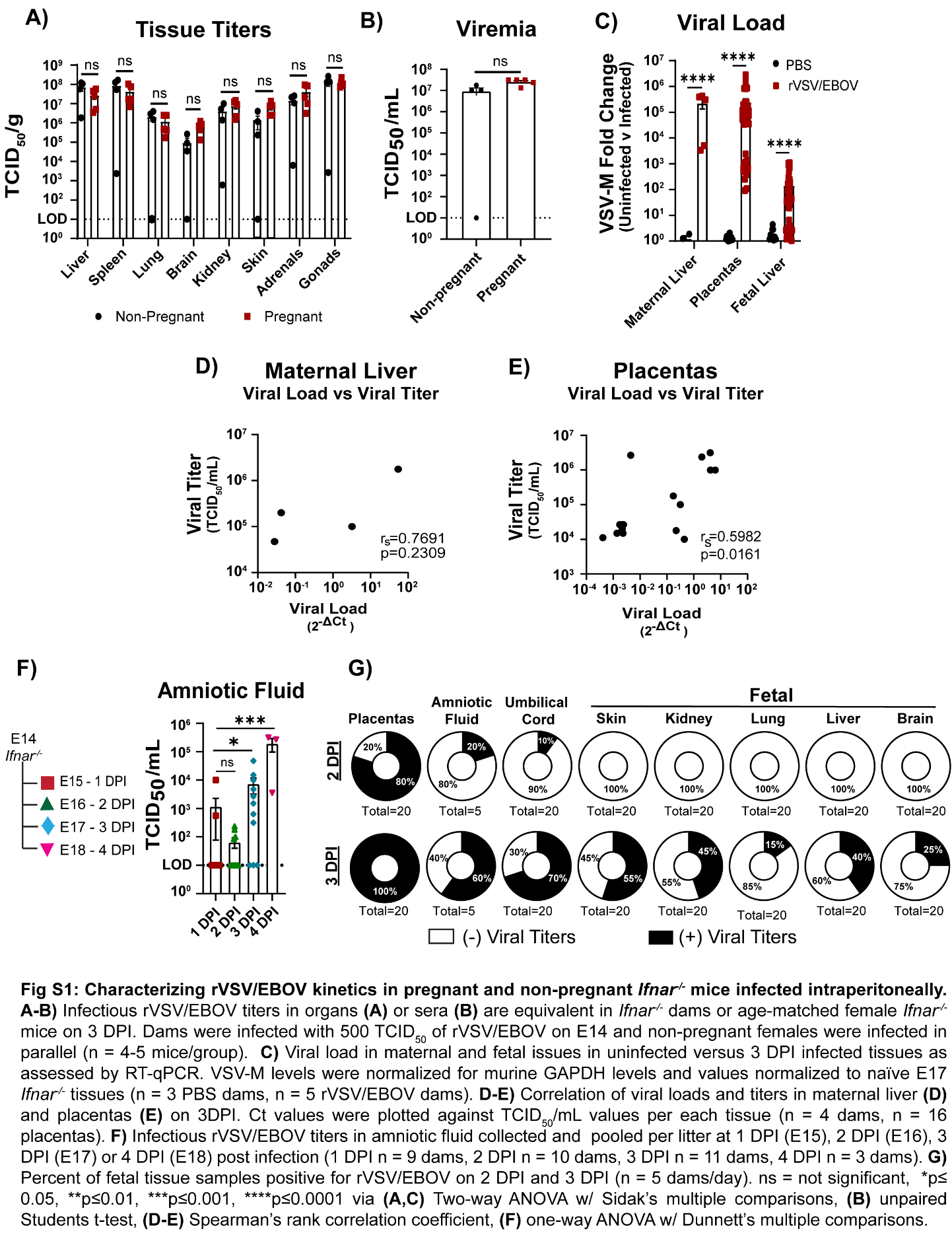

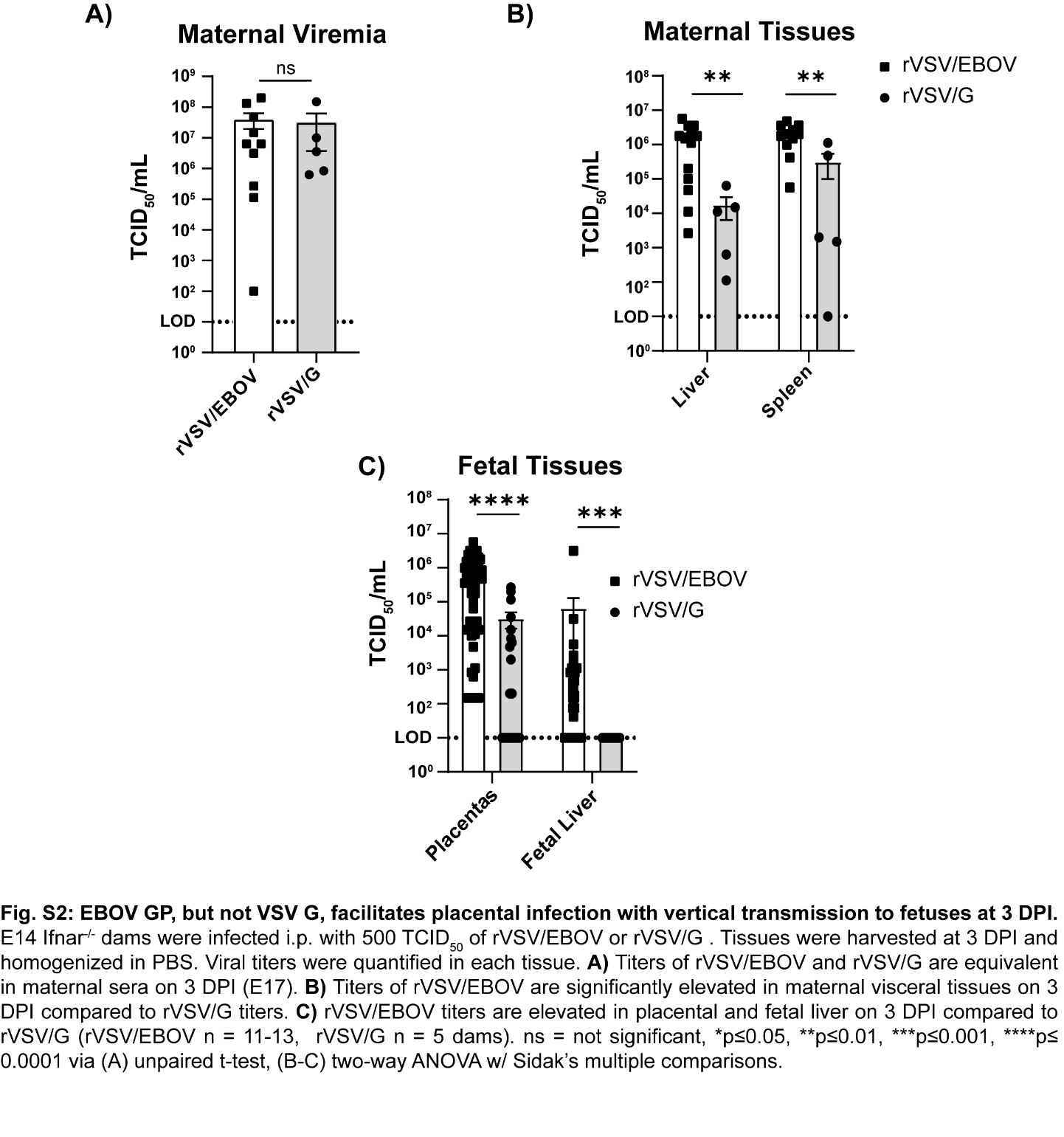

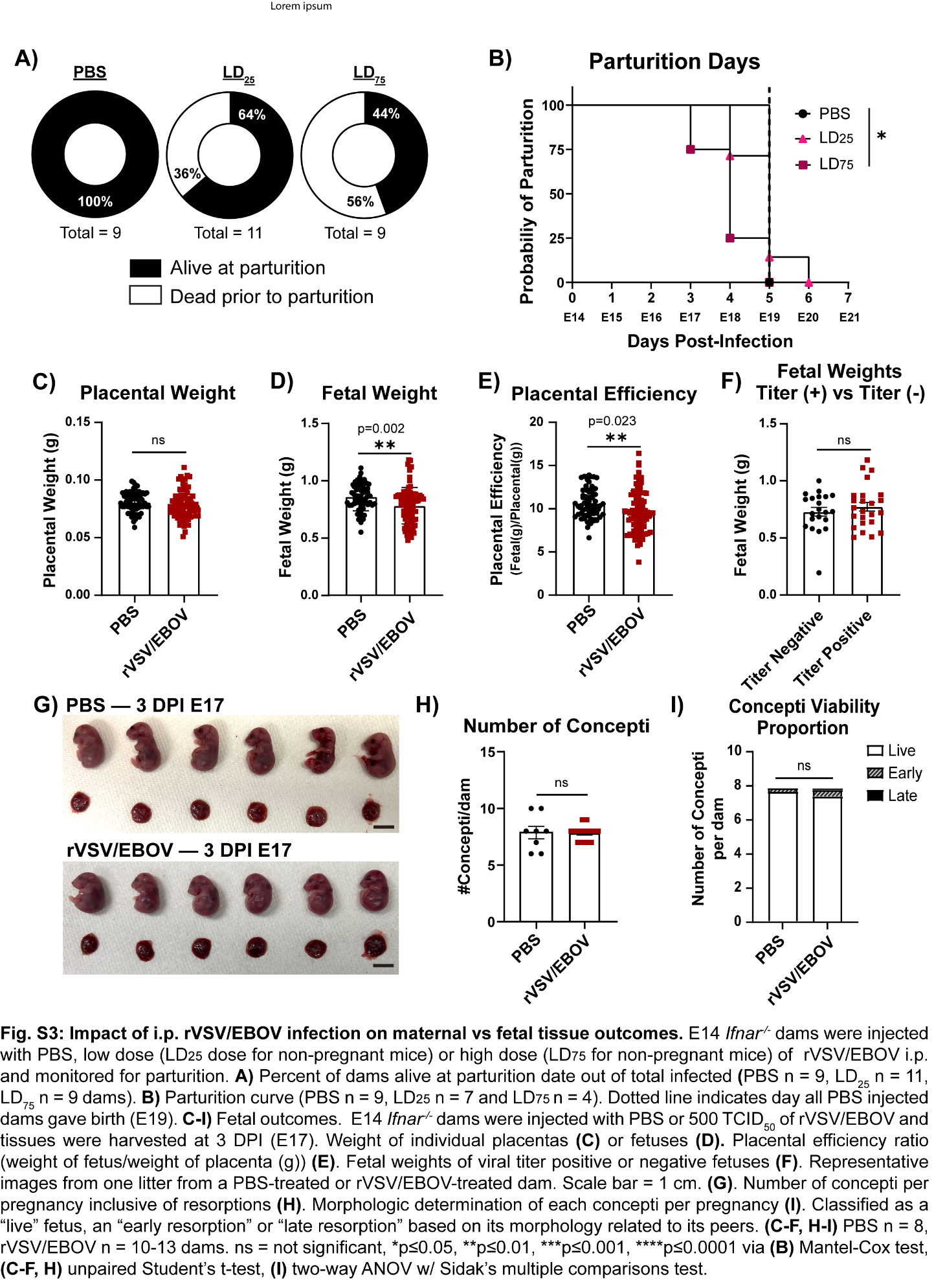

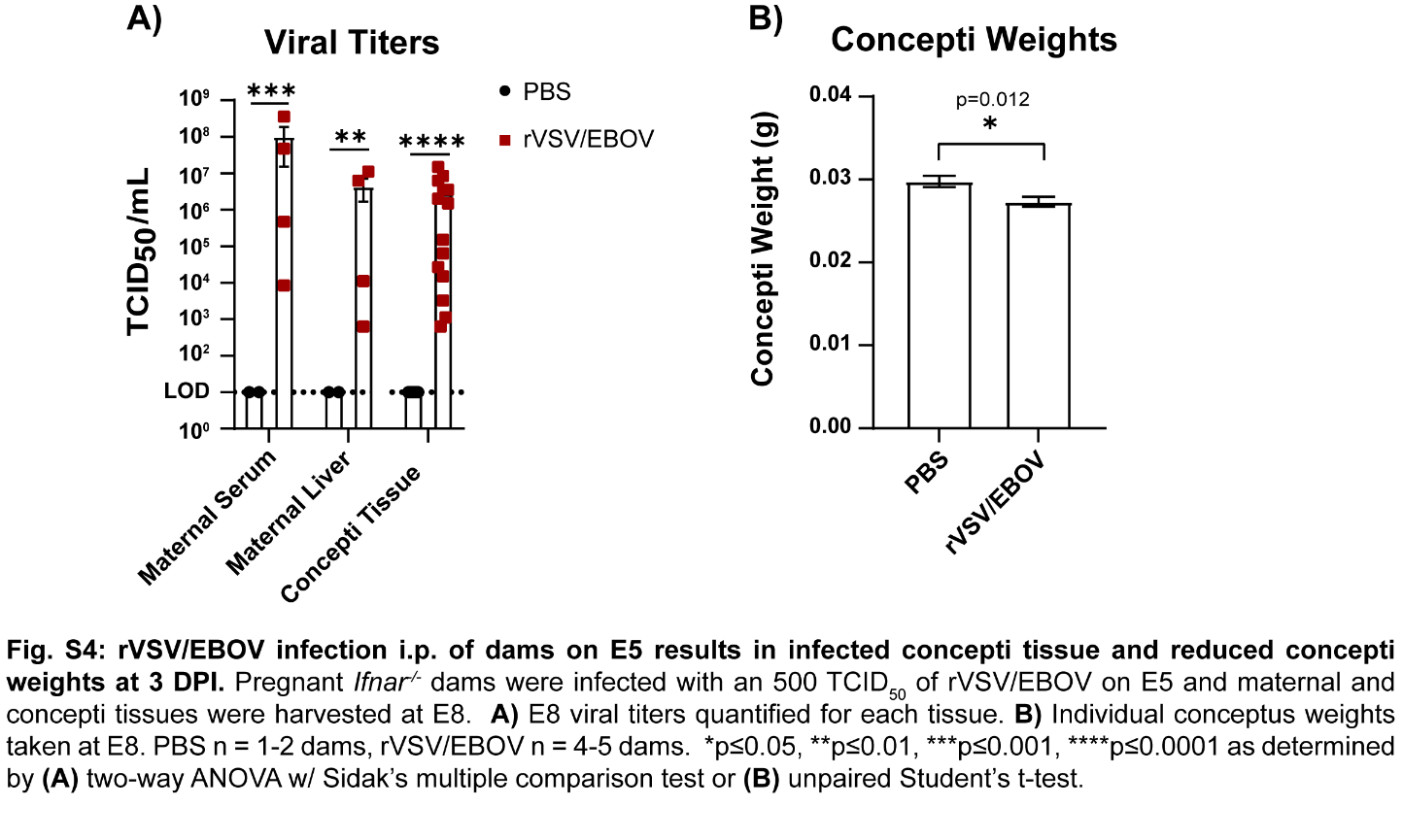

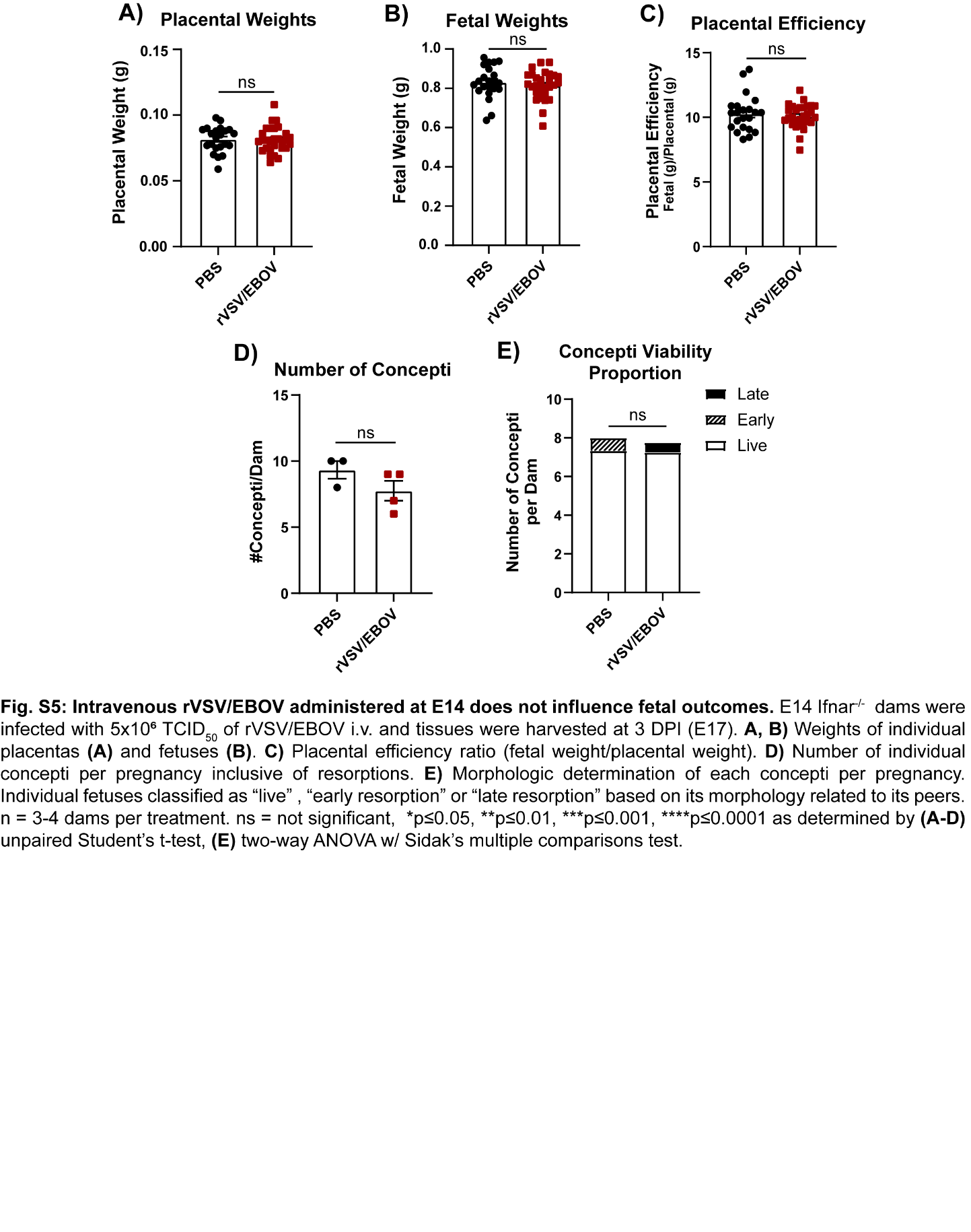

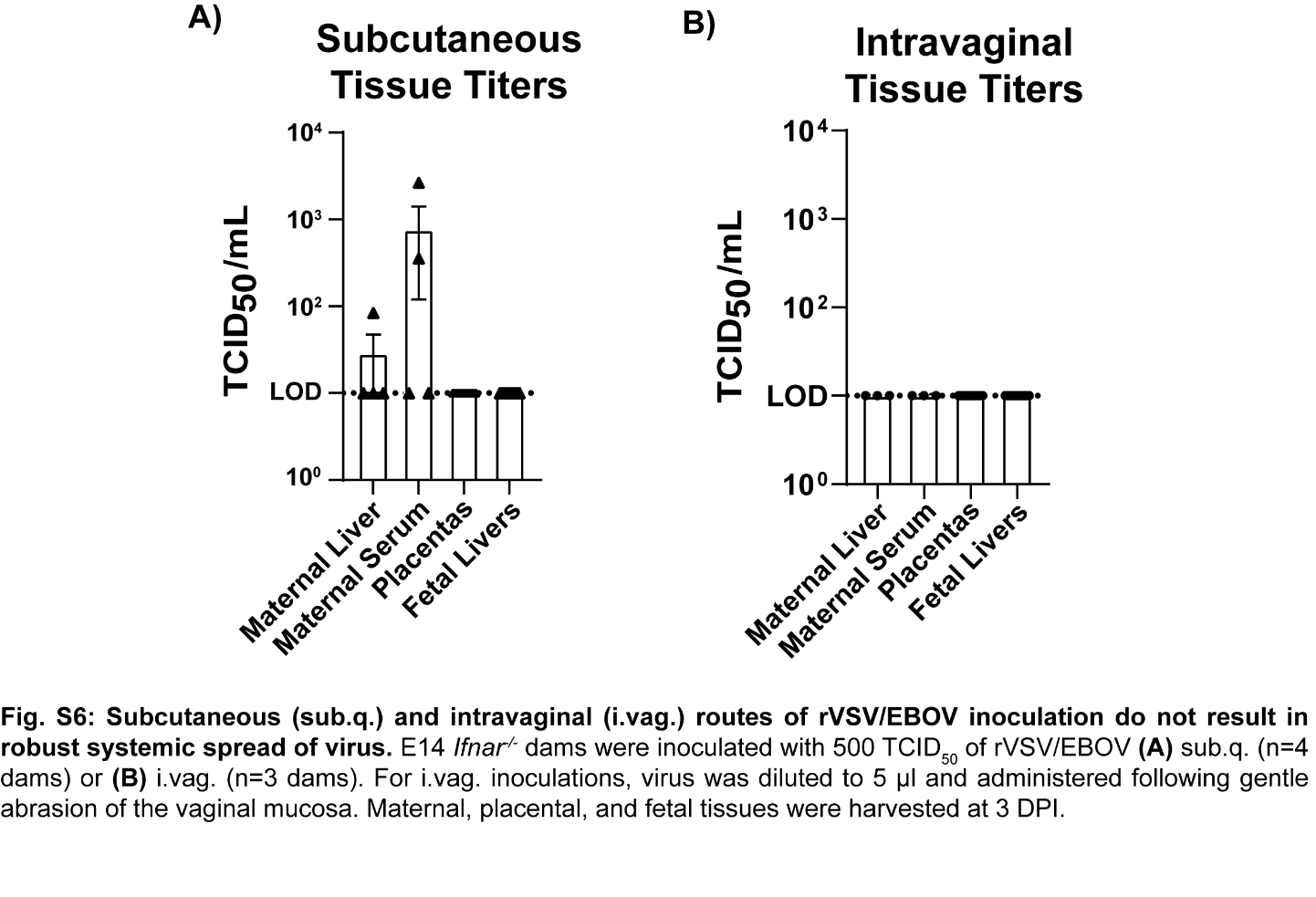

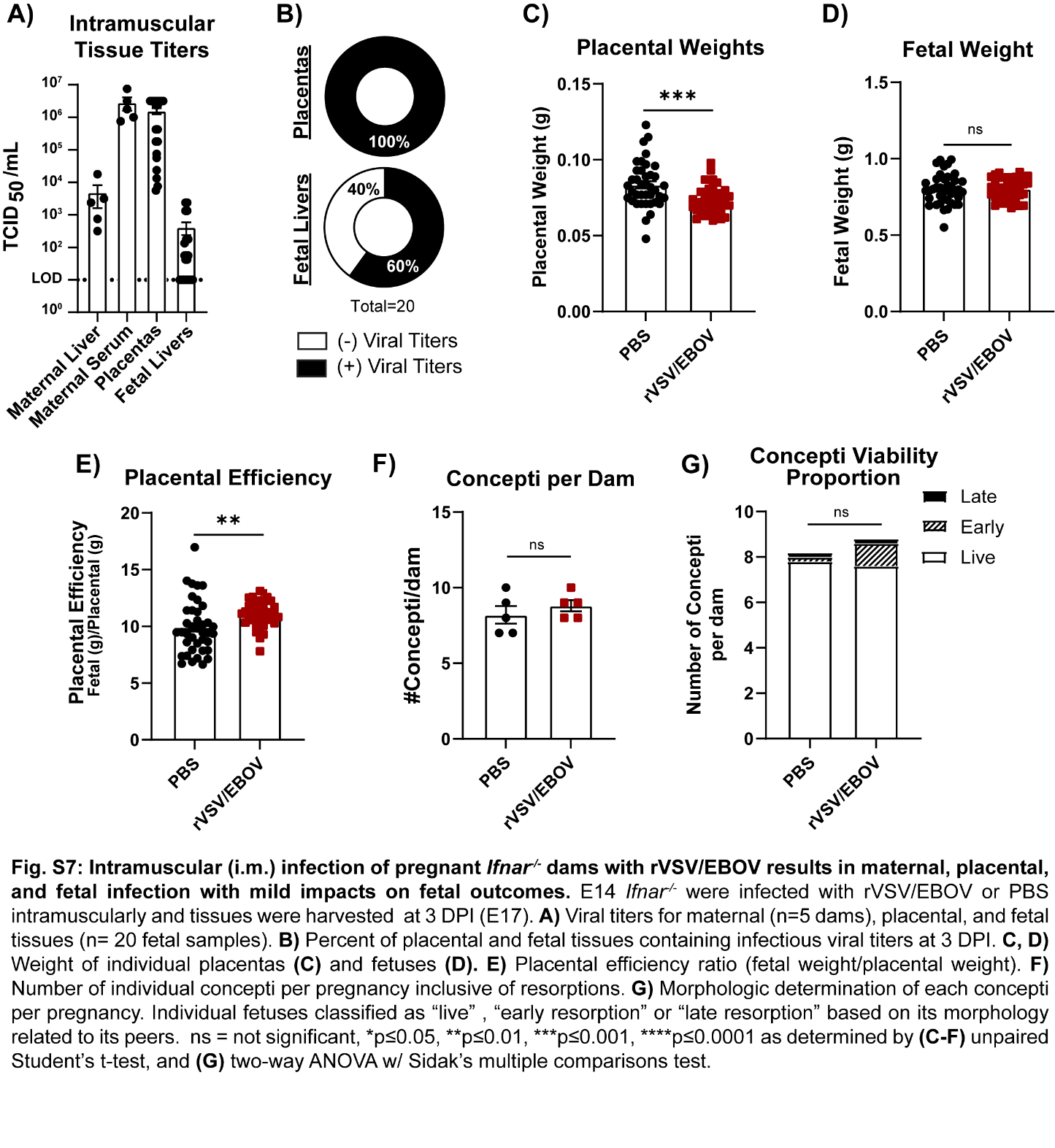

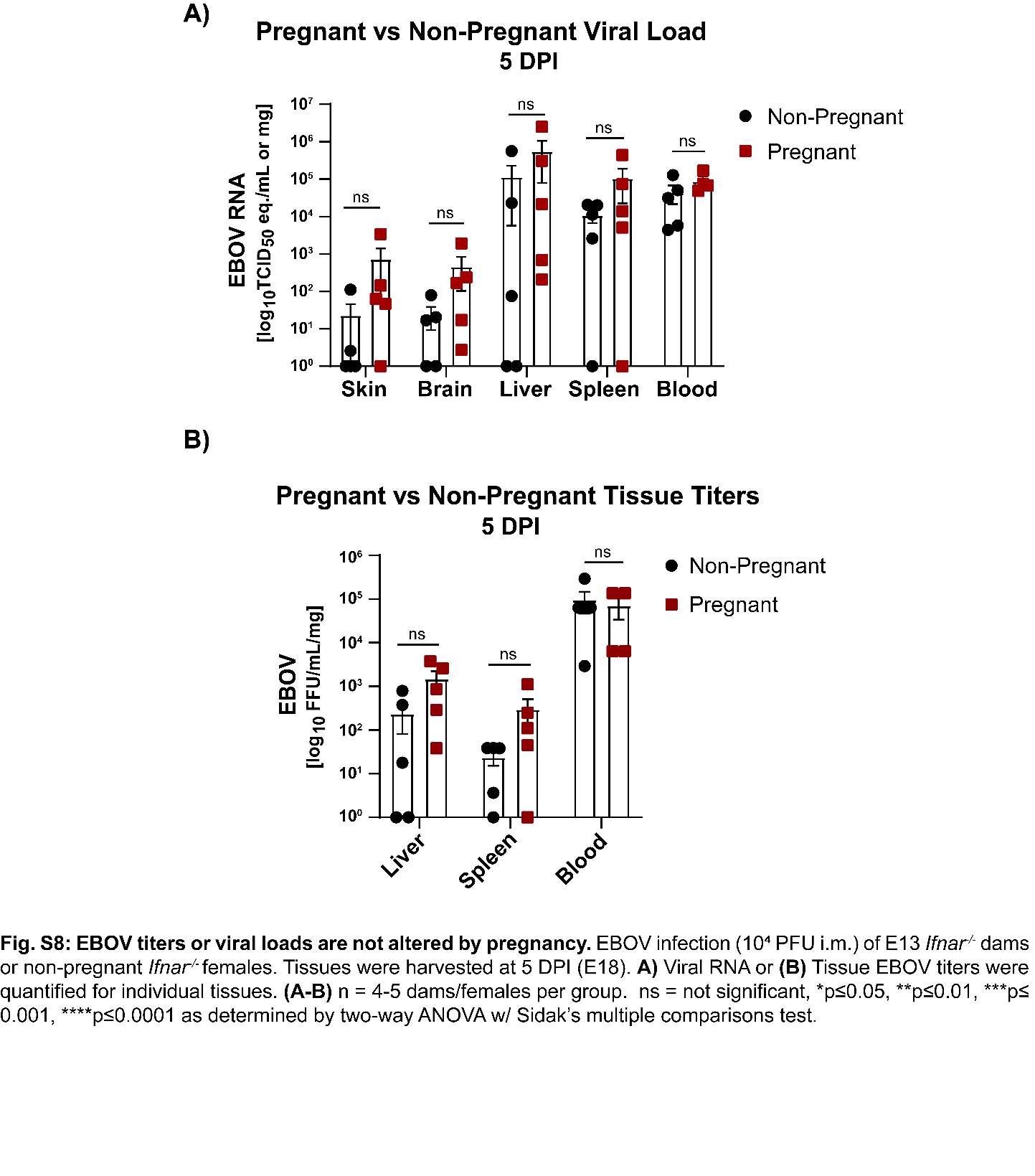

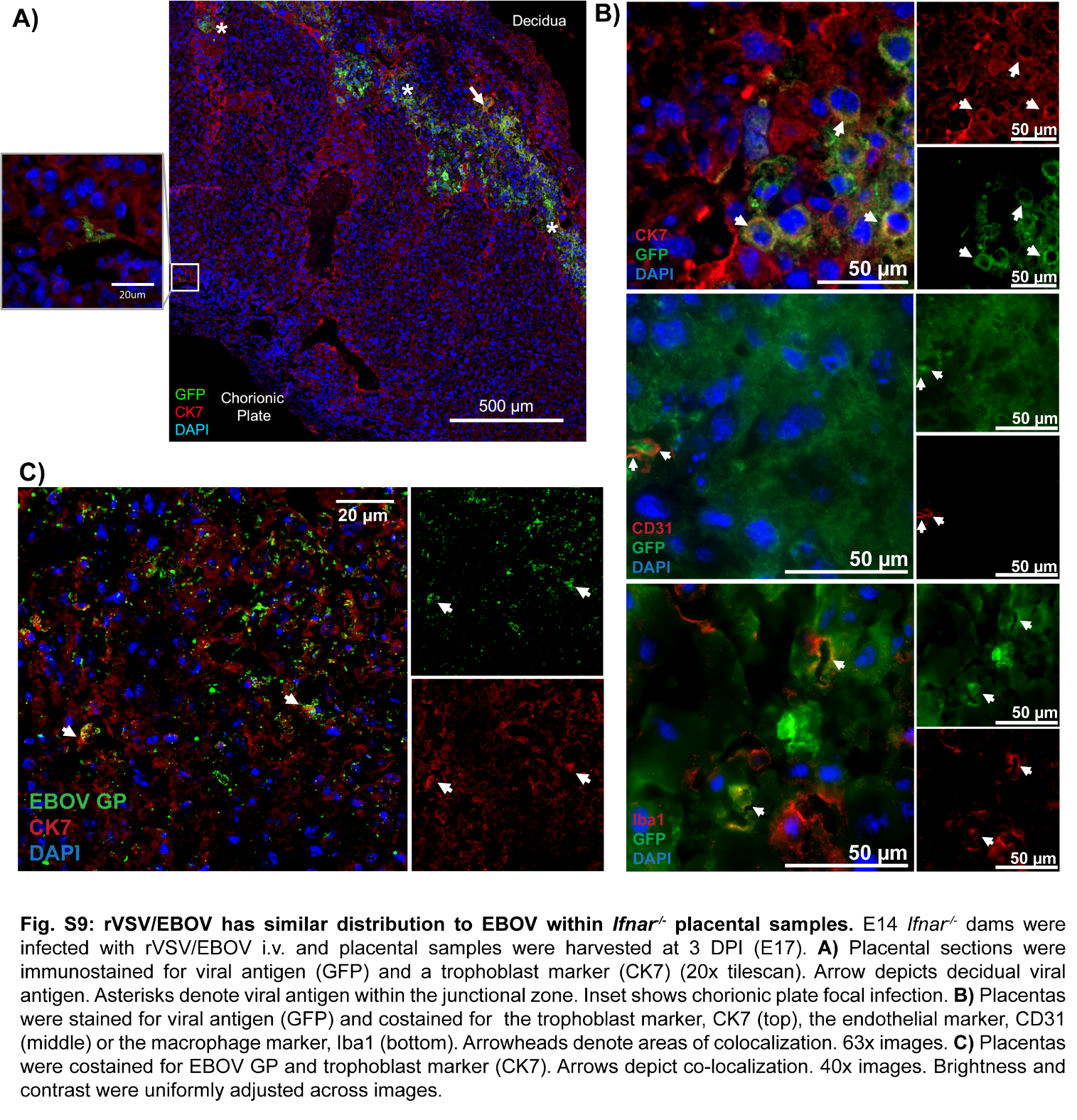

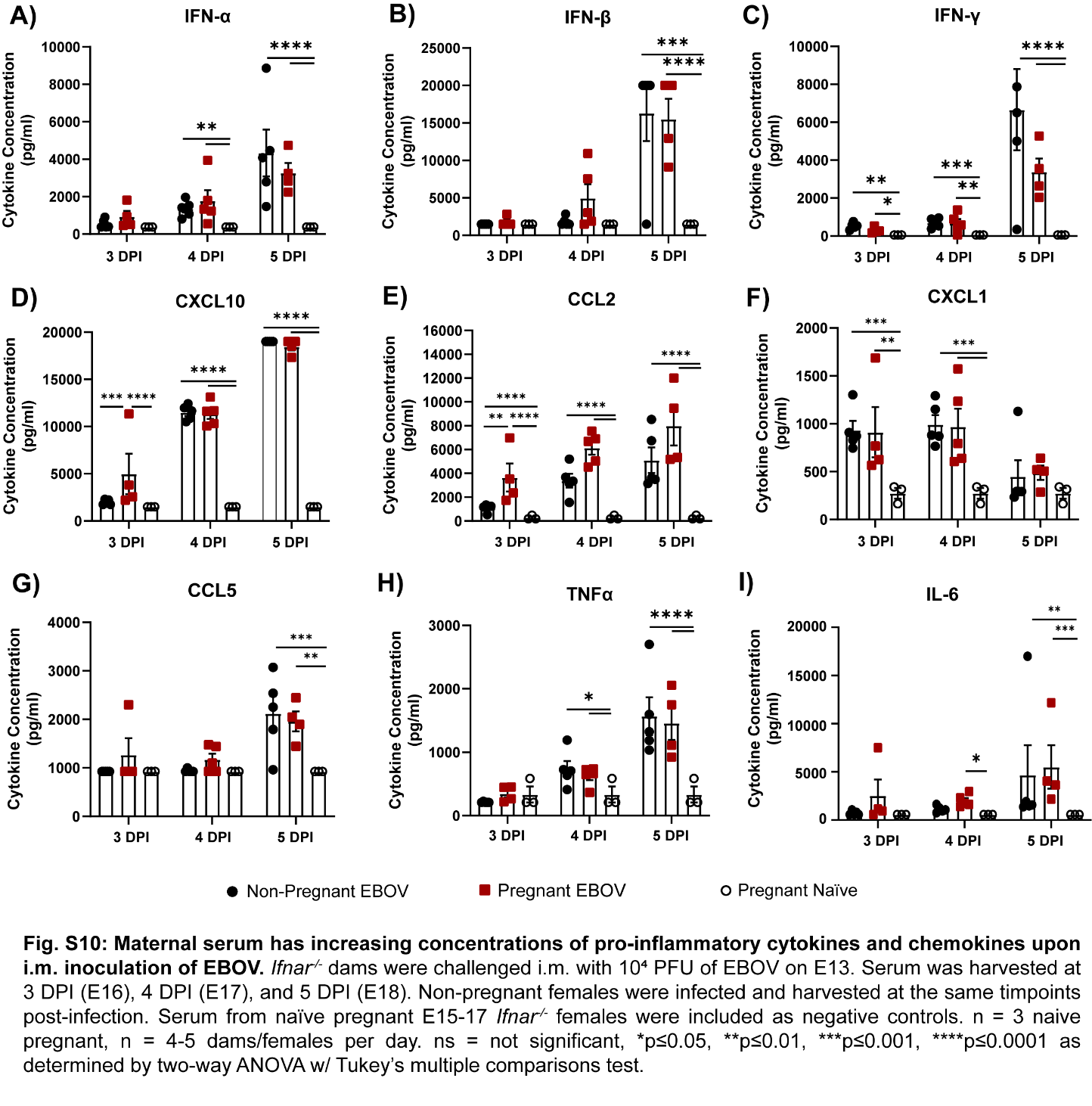

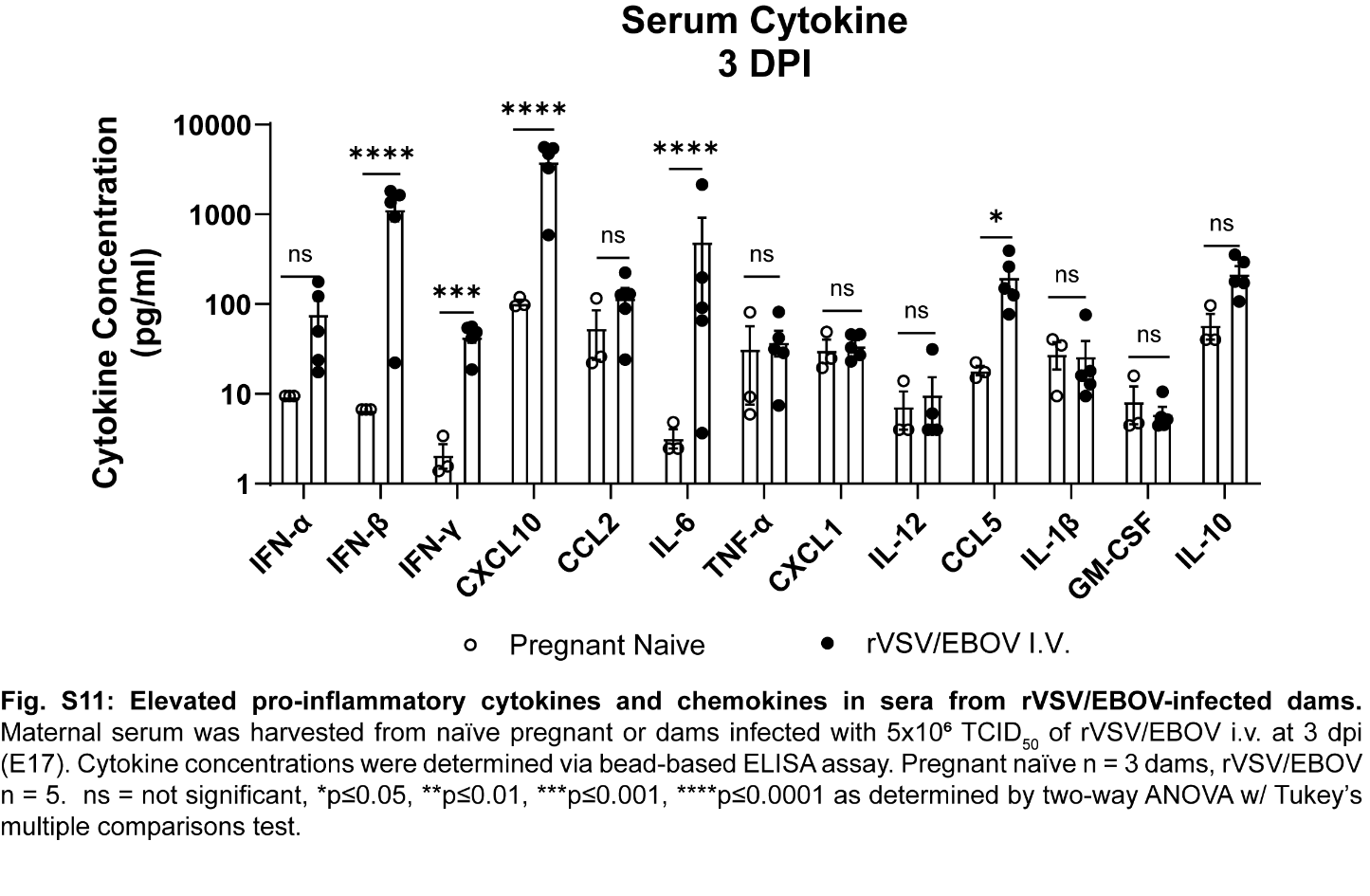

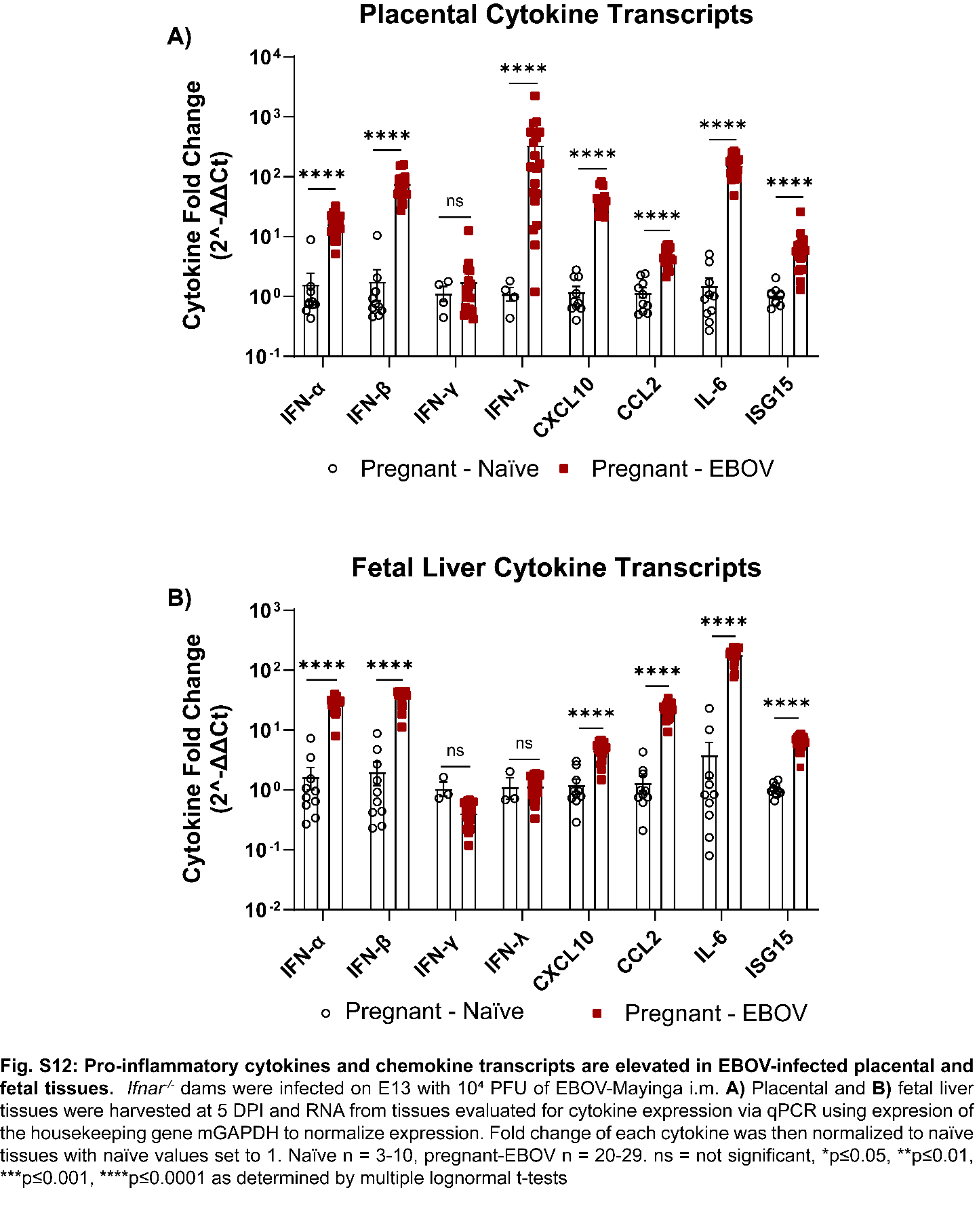

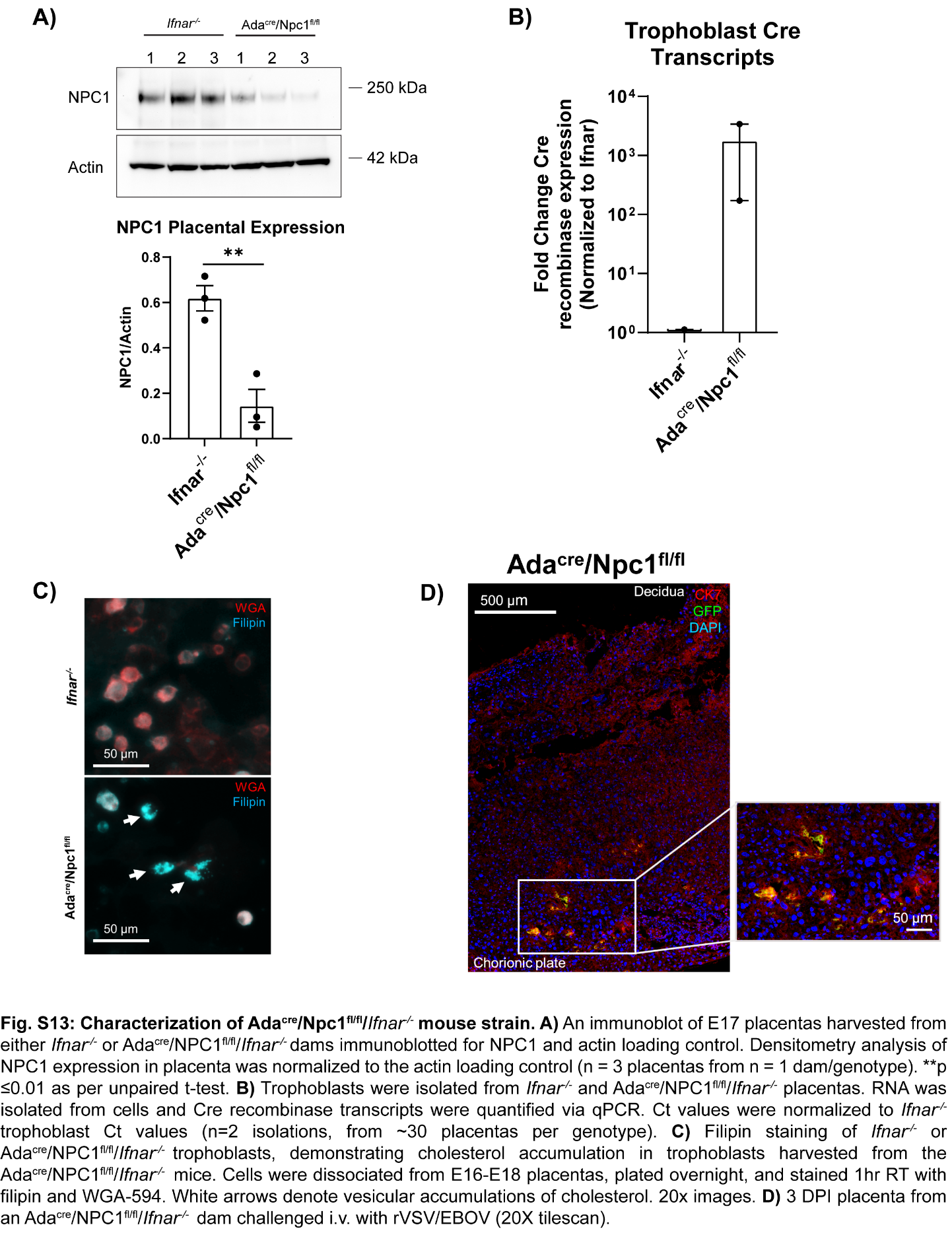

**Table S1: Murine genotyping primer sequences.**

| **Targeted Gene** | **Desired Final Genotype** | **Primers** |
| --- | --- | --- |
| Npc1 | Npc1^fl/fl^ | P1 – TACTTGGTAGTTGTCAGGTAGGCTTATGCT  P2 - GTCCACAGAACGGGTCATCT  P3 - ACACTGCAACGGGCTCCTTTG |
| Ada/Tpbpa | Ada^cre^ | F – CGG TCTCTGAGAGCCATC  R - CAGGTTCTTGCGAACCTCAT |
| Ifnar | Ifnar^-/-^ | Common F- CGA GGC GAA GTG GTT AAA AG  WT R – ACG GAT CAA CCT CAT TCC AC  Mutant R - AAT TCG CCA ATG ACA AGA CG |

**Table S2. qPCR primer sequences**

| **Gene Target** | **Primer Sequences (5’-3’)** |
| --- | --- |
| mGapdh | F – CAT CAC TGC CAC CCA GAA GAC TG  R – ATG CCA GTG AGC TTC CCG TTC AG |
| mIFNα | F – CCT GAG AGA GAA GAA ACA CAG CC  R – TCT GCT CTG ACC ACT TCC CAG |
| mIFNβ | F – AGC TCC AAG AAA GGA CGA ACA  R – GCC CTG TAG GTG AGG TTG ATC T |
| mIFNγ | F – CAG CAA CAG CCA GGC GAA AAA GG  R – TTT CCG CTT CCT GAG GCT GGA T |
| mIFNλ | PrimeTime^TM^ Mm.PT.58.8956530 |
| mIFNε | F – GAA ACG GAT TCC CTT CCA AT  R – ACT GCT GGA CTG ACG AGC TT |
| mCXCL10 | F – ATC CCT CTC GCA AGG ACG GT  R – CGG ATT CAG ACA TCT CTG CT |
| mCCL2 | F – TTA AAA ACC TGG ATC GGA ACC AA  R – GCA TTA GCT TCA GAT TTA CGG GT |
| mISG15 | F – TCT GAC TGT GAG AGC AAG CAG  R – ACC TTT AGG TCC CAG GCC ATT |
| mIL-6 | F – GAC TGG GGA TGT CTG TAG CTC  R – CAA CTG GAT GGA AGT CTC TTG C |
| Cre | F -- CCC TGT TTC ACT ATC CAG GT R – GGG TAA CTA AAC TGG TCG AG |
| VSV-M | F – CCT GGA TTC TAT CAG CCA CTT C  R – TTG TTC GAG AGG CTG GAA TTA G |

**Table S3: Antibodies, optimal dilutions, and incubation conditions for immunofluorescence**

| **Antibody** | **Dilution** | **Incubation** |
| --- | --- | --- |
| Human anti-EBOV GP (KZ52)  IBT Bioservices #0260-001 | 1:100 | 4ºC overnight |
| Rabbit anti-GFP Alexa 488  Invitrogen #A-21311 | 1:50 | 4ºC overnight |
| Goat anti-CD31  R&D #AF3628 | 1:50 | 4ºC overnight |
| Rabbit anti-mCK7  Invitrogen #PA5-29033 | 1:100 | 4ºC overnight |
| Rabbit anti-Iba1  Fujifilm #019-19741 | 1:100 | 4ºC overnight |
| Rabbit anti-F4/80  Cell Signaling #30325 | 1:100 | 37°C overnight |
